# Image Analysis Tools for Scanning Electron Microscopy

**DOI:** 10.64898/2026.03.11.711125

**Authors:** David H. Shtengel, Gleb Shtengel, C. Shan Xu, Harald F. Hess

**Affiliations:** University of Virginia, College of Arts & Sciences, Charlottesville, VA 22904, USA; Yale University, Cellular & Molecular Physiology, New Haven, CT 06520, USA; HHMI Janelia Research Campus, Ashburn, VA 20147, USA

## Abstract

Scanning Electron Microscopy (SEM) is widely used in many scientific fields, particularly in life sciences, offering high-resolution information on the ultrastructure of biological organisms. Accurate characterization of SEM image quality is important for assessing the SEM tool performance, in addition to sample preparation protocol, imaging conditions, etc.

This paper provides an overview of tools we developed as plugins for the popular image processing package Fiji (ImageJ) (*1*). These tools include signal-to-noise ratio analysis, contrast evaluation, and resolution analysis, as well as the capability to import images acquired on custom FIB-SEM instruments (*2*). We have also made these tools available in Python, with both versions available on GitHub.

## Introduction

An ability to accurately estimate the signal-to-noise ratio (SNR) and contrast in SEM images is important for many reasons. Determining the SNR on a known standard sample is a useful tool in assessing the quality of the SEM system alignment and performance. SNR and contrast information are also valuable in evaluating sample preparation, for example, the quality of staining. An accurate SNR metric is important to establish optimal imaging conditions across multiple samples and imaging systems.

Finally, if SEM images from multiple detectors are acquired simultaneously, overall SNR could be increased if the detector signals are added with appropriate weights. These weights should be proportional to the SNRs in the channels, so an accurate procedure for estimating the SNR is needed.

Resolution is another important metric for assessing the quality of the images and imaging systems. Accurate determination of resolution is important in making sure the imaging system performance is optimal and for properly setting the acquisition parameters such as pixel size, SEM current, landing energy, etc.

## Evaluation of the Signal-to-Noise Ratio

The image intensity can be expressed as a sum of signal *s* and noise *n* terms:

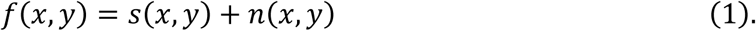

There are various ways to determine the SNR from images; SNR definitions also vary. We will follow papers (*3–6*), which define SNR as:

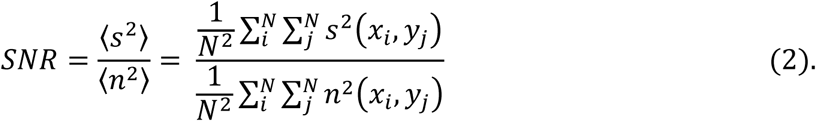

Multiple perfectly aligned images of the same area need to be collected in order to determine SNR from Eq (2) directly. This is usually impractical, and alternative methods for experimental evaluation of SNR have been proposed. Cross-correlation approach (*3*) allows determining SNR from just two images. However, its accuracy is degraded by the parasitic artifacts, such as line jitter, often present in SEM systems. Additionally, this method does require precise alignment of two images. Finally, it requires the knowledge of dark count, which is often missing if the images are saved with arbitrary offset (for example, in order to maximize the dynamic range of the images).

A single-image method based on the analysis of autocorrelation function (ACF) was proposed to address the above limitations (*6*). It lifts the requirement of two images and precise registration. However, it has some disadvantages as well. Estimation of noise-free autocorrelation is a critical and non-trivial step in the procedure. Multiple approaches have been proposed (*7*), but all have limitations, particularly if the pixel size is close to the image resolution. And this method also requires knowledge of dark count, which may not be readily available.

It should be noted that both cross-correlation and auto-correlation methods rely on the assumption that since signal and noise are uncorrelated their cross-product terms vanish in the analysis. This assumption is not correct in most general case and was demonstrated to be particularly erroneous in defining fixed Fourier Shell Correlation (FSC) thresholds and treatment of spectral SNR (*8*, *9*).

We propose a technique that allows determining the SNR from a single image without knowledge of dark count. In fact, our method allows determining dark count along with SNR.

In an SEM system, we have access to the signal from a detector that can be assumed to be linearly proportional to the number of detected electrons. The detector signal 𝑓*_d_* can be written as:

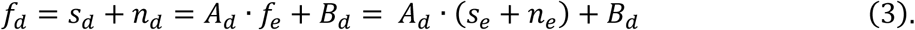

where 𝑠*_e_* and 𝑛*_e_* are electron signal and noise, 𝑠*_d_* and 𝑛*_d_* are the detector signal and noise, and 𝐴*_d_* and 𝐵*_d_* are the detector scaling coefficients (additive detector noise is neglected for now). The detector scaling coefficients are not known; determining them is desirable.

We can write the following:

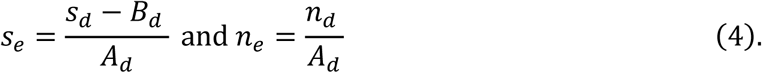

Then SNR can be written as:

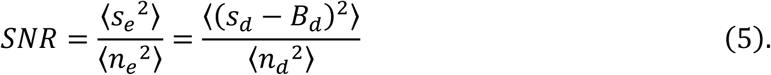

So, we only need to know the offset 𝐵*_d_* to determine SNR, but it turns out that we can easily determine both 𝐴*_d_* and 𝐵*_d_* using the following argument. In the case of the Poisson process, the variance and signal should be equal, 〈𝑛*_e_*^2^〉 = 𝑠*_e_*. Therefore, the measured detector intensity and variance should be scaled as:

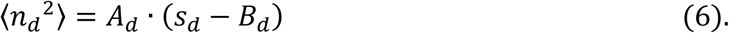

If the sample can be measured repeatedly, variance vs intensity can be analyzed using Eqs. (5) and (6), yielding the values of SNR and detector scaling coefficients 𝐴*_d_* and 𝐵*_d_*. If repetitive measurements are impractical, a modified approach using the same underlying math can be implemented. If the pixel size is smaller than the imaging system resolution, then smoothing the image and subtracting the smoothed data from the original data would provide both intensity (smoothed image pixel value) and variance (squared difference between the original and smoothed pixel values). Therefore, a simulated variance vs. intensity dependence can be built from a single measurement.

The requirement that the pixel size be smaller than the imaging system resolution is an important limitation. However, even if the pixel size is close to the resolution value, the above algorithm can be used with an additional step. We can calculate the image intensity gradient map and then exclude the pixels with high local gradients, effectively reducing the resolution of the remaining data set.

The process is illustrated in Figure 1. The original image (top left) is a signal from the secondary electron detector collected from a standard Sn-on-C (Tin on Carbon) SEM calibration sample (Ted Pella). It is then smoothed, and the difference is calculated. The pixels with very low and very high local intensity are excluded from subsequent analysis (pixels marked red and cyan, respectively, in the top-right image). Also, the pixels with high local intensity gradients (pixels marked blue in the top-right image) are excluded from subsequent analysis. The dependence of squared difference on smoothed intensity is then analyzed, yielding the values of SNR and detector dark count 𝐼_0_, as shown in Figure 1 (bottom).

**Figure 1.**
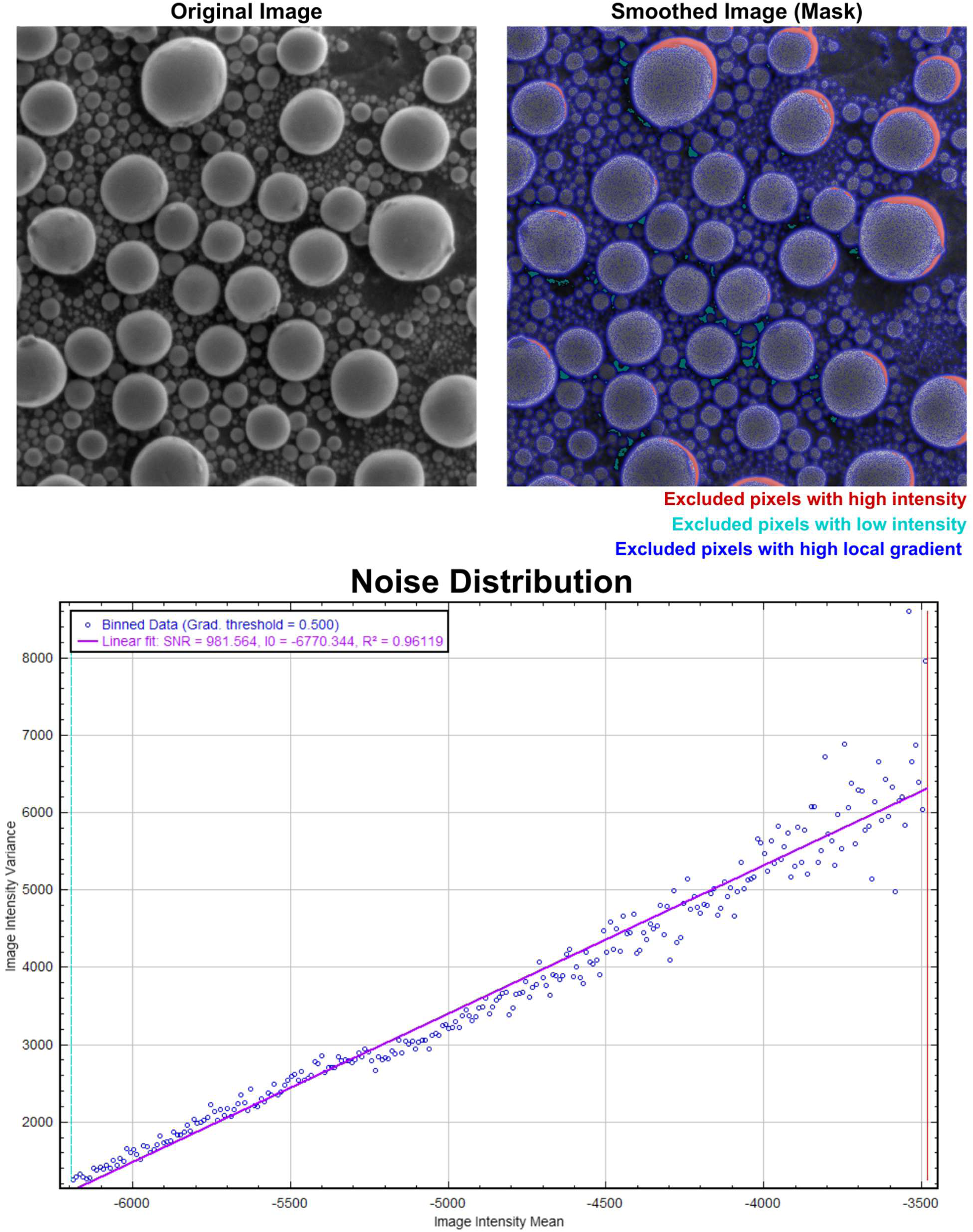
Analysis of the simulated Variance vs. Mean dependence, used to determine SNR and dark count (zero variance intercept).

The effect of the setting of the value of the gradient threshold on the analysis results is illustrated in SI Figure 6. When all pixels are included in the analysis (gradient threshold set to 1.0), the variance vs. intensity curve is not straight because the intensity variance at points with high local gradients is higher than it would be if it were solely determined by shot noise. As the value of the gradient threshold decreases, the variance vs. intensity curve becomes more linear, and the values of SNR and detector dark count stop changing. At very low values of gradient threshold, a majority of the pixels are excluded, and the analysis becomes less accurate. The setting of the gradient threshold between 0.25 and 0.5 generally provides accurate results.

It is important to note that this approach works only if the detector is linear and no nonlinear image intensity post-processing (such as, for example, CLAHE, (*10*)) was performed.

Also, averaging or band-limiting the data acquisition, as well as image transformations that involve registration (e.g., stack registration), include interpolation steps that will affect the SNR value and may affect the determination of the detector dark count 𝐼_0_.

## Evaluation of Image Contrast

Another important parameter of the SEM image is image contrast, a ratio of the data range to the average value of the data. Along with SNR, contrast characterizes the quality of heavy metal staining in the SEM samples. While SNR is proportional to the average level of staining, the contrast characterizes the specificity and selectivity of staining. In an ideal sample, both SNR and contrast should be as high as possible.

We can define image contrast as:

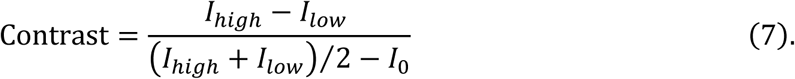

where 𝐼*_high_* is the average signal in highly stained areas, such as membranes; 𝐼*_low_* is the signal in areas with low stain, such as intercellular space; and 𝐼_0_ is the detector dark count. Knowledge of all three values: 𝐼*_high_*, 𝐼*_low_*, and 𝐼_0_ is required to use Eq. (7). Dark count 𝐼_0_ can be determined from the variance vs. intensity curve as described in the previous section. 𝐼*_high_* and 𝐼*_low_* can be determined directly from the probability distribution function (PDF). The PDF of the image intensity of the sample in Figure 1 is shown in Figure 2.

**Figure 2.**
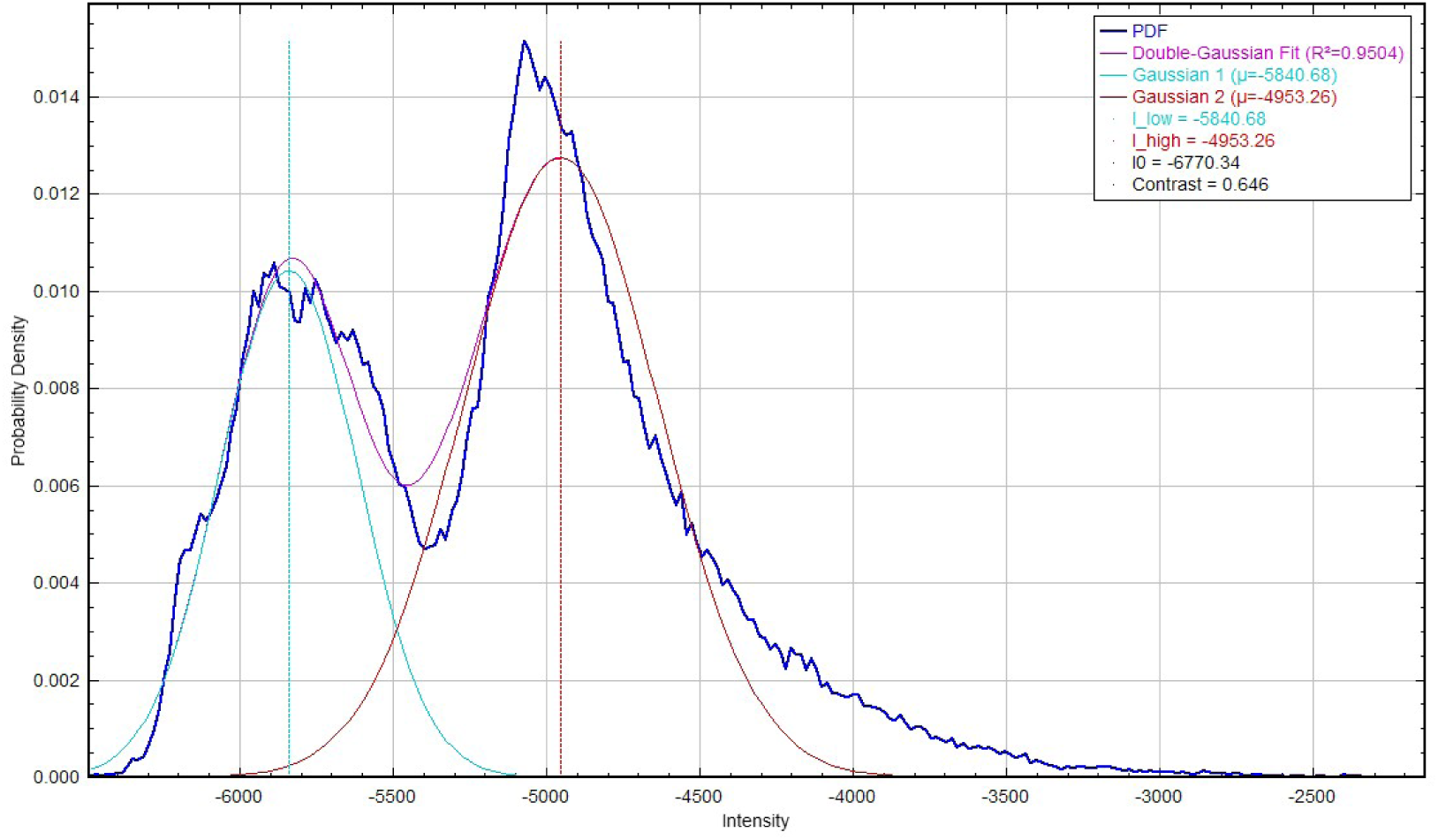
PDF of the image intensity of the sample in Figure 1, illustrating determination of I-high and I-low values and the value of image contrast.

The two peaks in the PDF of the SEM image of the Sn-on-C sample in Figure 2 correspond to the signals from the tin surface (𝐼*_high_*) and carbon surface (𝐼*_low_*). In the case when two PDF peaks 𝐼*_high_* and 𝐼*_low_* have similar counts, they correspond to 75% and 25% levels in the cumulative distribution function (CDF). The intensities corresponding to these CDF levels can be used as an alternative way to define and determine contrast in the case that the PDF does not have a clear bimodal structure. Similarly to the noise analysis, the evaluation of the contrast is accurate only if no non-linear image intensity transformation was performed.

It should be noted that staining levels vary between different parts of cells and tissue. Relative spatial densities of these differently stained parts may also vary. All of these affect the results of the contrast analysis. This is illustrated on the example of cultured cell, see SI Figure 13 - SI Figure 18 and SI Table 1. The staining level of ribosomes and chromatin are much higher than that of lipid membranes. Depending on the relative fraction of the ROI, covering the highly stained chromatin, the value of contrast determined by automatic double-Gaussian fit varies from 0.058 (yellow rectangle ROI, SI Figure 17) to 0.093 (cyan rectangle ROI, SI Figure 13) to 0.171 (red rectangle ROI, SI Figure 15). Moreover, if we select I*_low_* and I*_high_* levels manually to correspond to cytosol and membranes, the corresponding contrast is 0.188 (SI Table 1, middle row), and with I*_low_* and I*_high_* levels manually set to correspond to cytosol and ribosomes, respectively, the contrast is 0.325 (SI Table 1, bottom row).

In order to address the above uncertainties in the process of determining the image contrast, we developed GUI that allows switching between the automatic and manual selection of I*_low_* and I*_high_* levels and visualization of this selection (SI Table 1).This could be helpful in the process of optimizing the staining protocols targeting specific structures.

## Evaluation of Image Resolution

Resolution of the images can be evaluated directly by analyzing the transitions across the sharp features in the image or indirectly by analyzing the frequency representations of the SEM data. Multiple methods based on Fast Fourier Transforms have been proposed (*11–13*). These methods work well for TEM images, but not for SEM images due to the line-scanning nature of SEM and artifacts such as line jitter.

Direct analysis of sharp transitions is more practical for SEM images. The distance of 37% to 63% intensity change across a sharp edge is a metric often used for the image resolution of SEM systems.

We implemented this procedure, using the following processing steps:

1. Calculate the gradient map and create a list of potential transition points for evaluation. Sort these points in order of absolute values of the gradient.
2. Build a set of evaluation points using the following procedure:

2.1. Select a point with the maximum absolute value of the gradient and add it to the set.
2.2. Draw an exclusion circle around it and remove all points within that circle from further consideration.
2.3. Go to the next point and repeat.
3. For the evaluation points determined in the previous step, evaluate the edge transitions:

3.1. Use local intensity gradient to determine the transition direction.
3.2. Isolate a square subset of the image around the center of the evaluation point.
3.3. Build a section of the image (trace) along the transition direction using nearest neighbor interpolation and find min and max values of that trace. Analyze only the “good” transitions that have min and max values close to the min and max values of the entire image.
3.4. For all “good” transitions determined above, find their 37% and 63% transition distances using linear interpolation.

The results of this resolution analysis for the Sn-on-C sample used in Figure 1 are shown in Figure 3. The original image is shown in Figure 3, top left. The centers of analyzed traces are indicated with colored points. The individual transition plots are shown in Figure 3, bottom left (colors of the traces correspond to the point colors in Figure 3, top left). The distribution of 37% and 63% transition distances determined from these traces is shown in Figure 3, top right.

**Figure 3.**
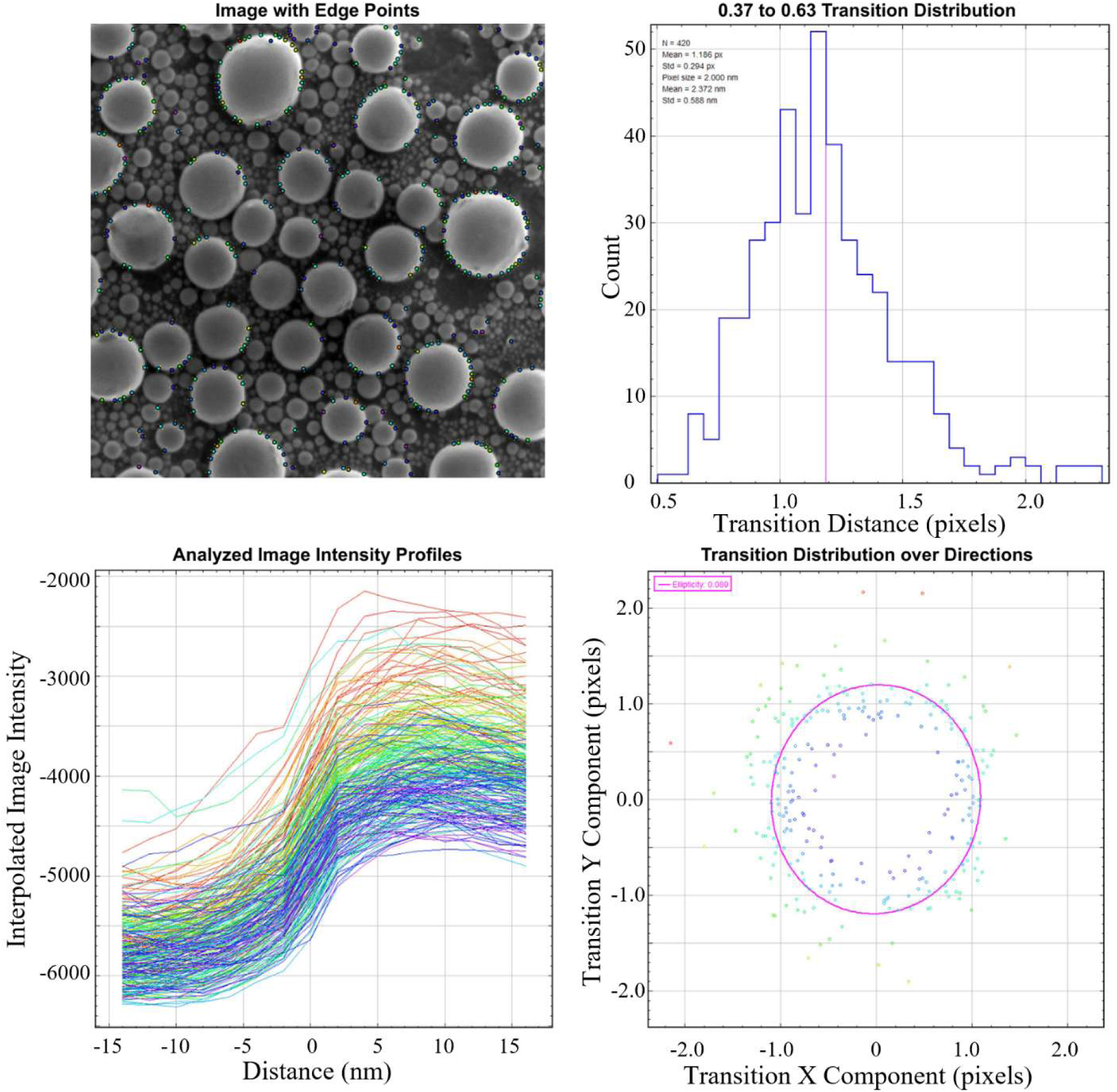
Analysis of the transitions across sharp edges to characterize the image resolution.

The 2D map of x- and y-components of the transitions (in Figure 3, bottom right) is used to determine ellipticity and is useful in characterizing the stigmation of the image or system.

## Results

We have performed SNR and contrast evaluation, the results are shown in Table 1 and SI Figure 11 - SI Figure 22. The values of SNR and Image Contrast vary over the range of selected samples. In addition, they may vary within a sample, depending on the selection of ROI, as illustrated on the example of Killer T-Cell attacking cancer cell (SI Figure 14 and SI Figure 16). It is important to keep this in mind when using these tools to compare different samples.

**Table 1.**
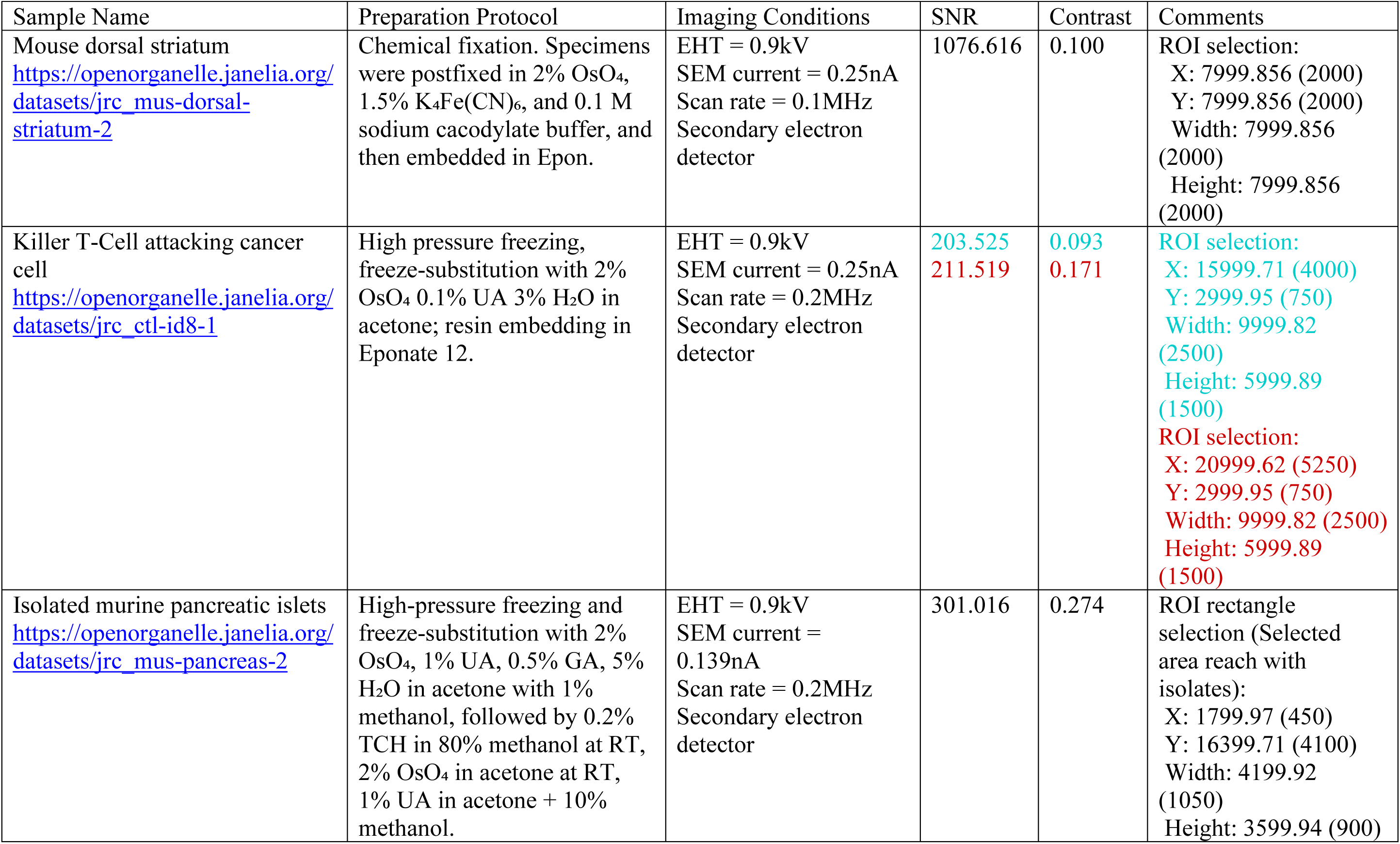
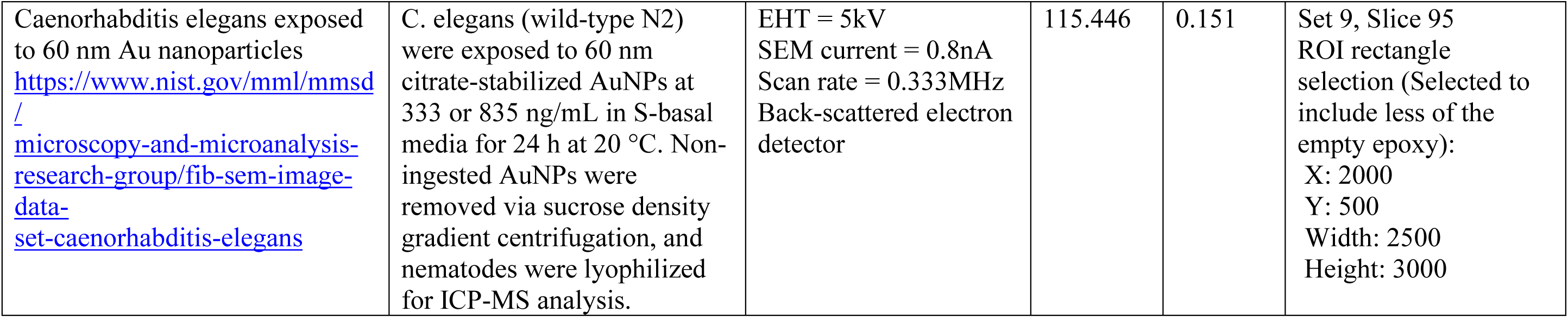
Results of the SNR and Contrast Analysis on select published datasets.

## Summary and Discussion

We developed signal-to-noise ratio analysis, contrast evaluation, and resolution analysis as FiJi plugins. These procedures and metrics can be helpful in quantifying the difference between SEM datasets, which, in turn, should help in both optimizing the sample preparation protocol and evaluating the SEM instrument performance.

It should be noted that both SNR and imaging contrast may further depend on imaging conditions, such as primary electron dose, landing energy, working distance, sample bias, and detector configuration. In order to make meaningful conclusions about the sample preparation, we need to either compare the data acquired under similar conditions, or account for differences in the imaging conditions.

It is also important to note that the image contrast may also vary for different parts of ultrastructure. For example, for cultured cell the contrasts are different for mitochondria, ER, and ribosomes/chromatin.

We have demonstrated how these procedures can be used on various published FIB-SEM datasets.

The plugins are available on GitHub.

## Acknowledgements

We thank Ken Hayworth and David Peale for helpful discussions, and to Martin van Heel for pointing out problems in treating uncorrelated signals as orthogonal in SNR analysis.

## Contributions

HFH conceived the concept, DHS, CSX and GS wrote the code. All authors discussed the results and wrote the correspondence.

## Use of AI Tools

This work utilized Claude Opus 4.6 (Anthropic, 2025) to assist with minor code debugging during the plugin development.

## Supplemental Information

### Fiji plugin installation and use

- Install the Fiji plugin by copying the fib_sem-*.*.*.jar file into the Fiji plugin directory:
- Restart Fiji.
- Open the source image.
- Use Fiji plugins by navigating to Plugins → FIB-SEM → Noise Statistics Analysis, etc. as illustrated in SI Figure 1.

**SI Figure 1.**
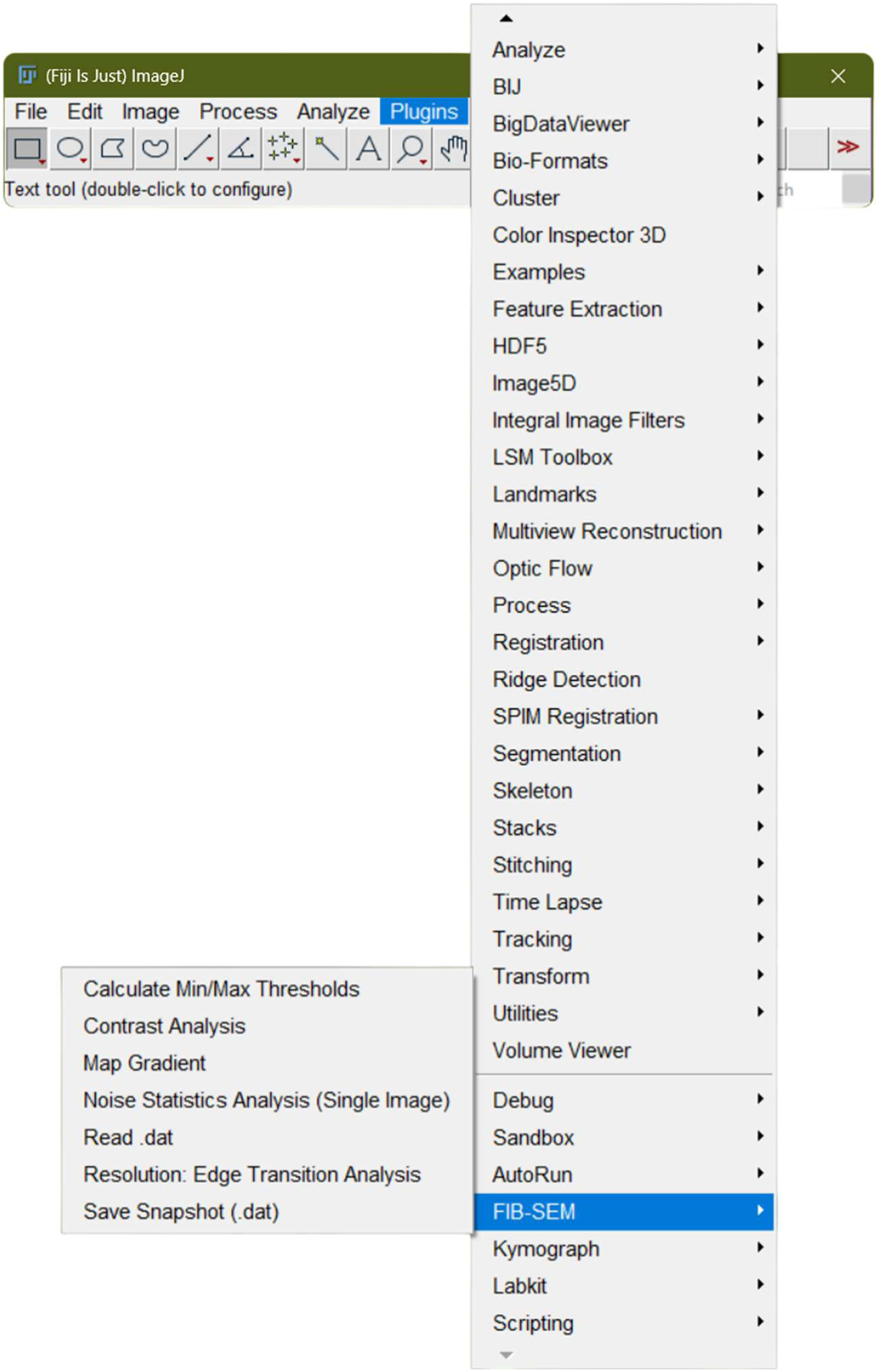
Using Fiji plugins.

### Importing the custom binary FIB-SEM data

The data files acquired on custom FIB-SEM instruments (*2*) contain important information in the headers. We created a plugin to import data. In order to import a custom .dat file, navigate to Plugins → FIB-SEM → Read .dat and enter the file name in the pop-up menu. The program will import the data and display the images. The header information can now be accessed by navigating to Image → Show Info, as illustrated in SI Figure 2.

**SI Figure 2.**
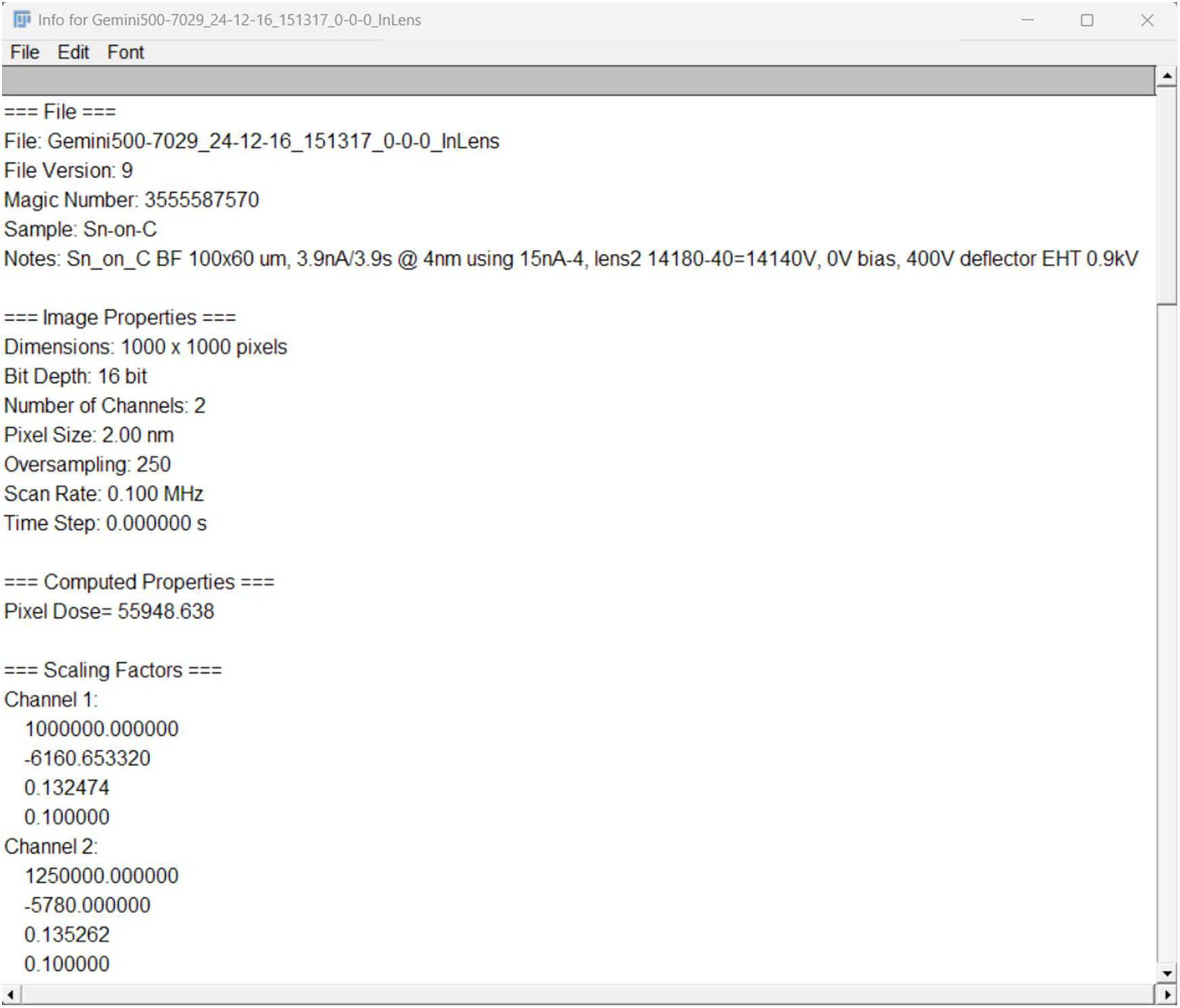
Header information displayed in the Image → Show Info window.

### Save a Snapshot of the FIB-SEM data

Once the image has been imported, a “snapshot” overview can be generated for quick reference. This snapshot is automatically saved to the same directory as the .dat it displays. Snapshot creation can be called by navigating to and clicking Plugins → FIB-SEM → Save Snapshot (dat), where there are no dialog options. An example snapshot will resemble the one illustrated in SI Figure 3. Example snapshot.

**SI Figure 3.**
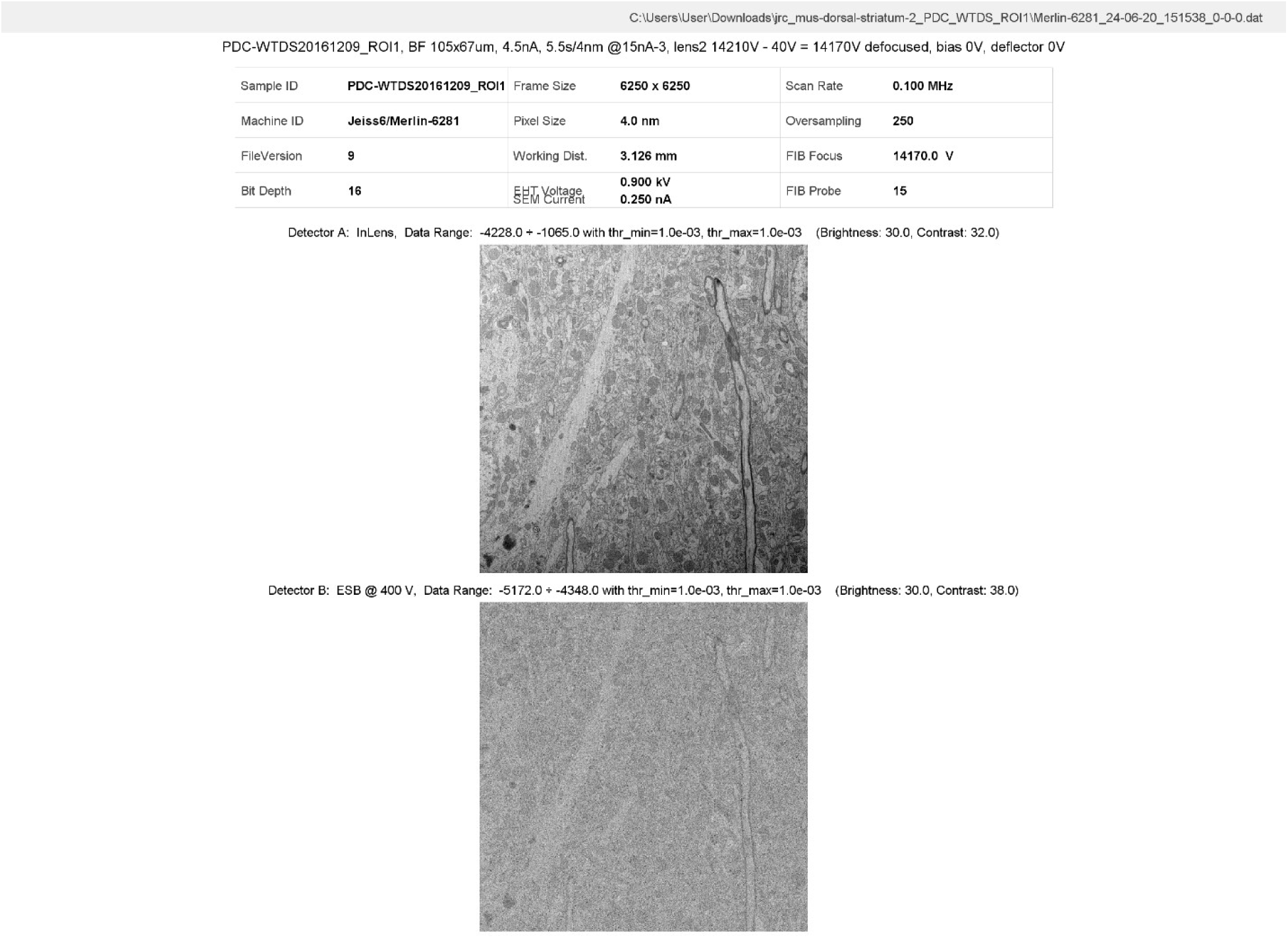
Example snapshot.

### Performing the Noise Analysis

Once the image has been imported (it can be any image, not just a custom .dat file), and a rectangular region of interest (ROI) has been defined, the noise analysis can be performed by navigating to Plugins → FIB-SEM → Noise Statistics Analysis (Single Image). The pop-up menu with settings for Noise analysis will appear as illustrated in SI Figure 4.

The parameters are:

**Analysis CDF threshold (lower)** – lower boundary of cumulative distribution function (CDF) of the data used for analysis. Default is 0.01, meaning that the pixels with values below 0.01 of the CDF will be ignored.

**Analysis CDF threshold (upper)** – upper boundary of cumulative distribution function (CDF) of the data used for analysis. Default is 0.01, meaning that the pixels with values above the 99.99^th^ percentile of the CDF will be ignored.

These two bounds help filter the data that may fall within the detector’s non-linear response range.

**Number of bins (analysis)** – number of bins used for analysis of the data. Default is 256.

**Gradient threshold (upper)** – upper boundary for the gradient threshold. This parameter is set to exclude the pixels with high local gradients, which may have higher intensity variations because of fast-varying signal intensity, not due to noise. The effect of the gradient threshold setting on noise analysis results is discussed in more detail in the section below. The default setting is 0.5, meaning that only 50% of the pixels with the lowest local intensity gradients are considered in the noise analysis.

**Dark Count (intensity at zero variance)** – This is used if the user wants to override the detector dark count 𝐼_0_ determined automatically and to calculate the SNR value (labeled as SNR1) with a different detector dark count value.

**SI Figure 4.**
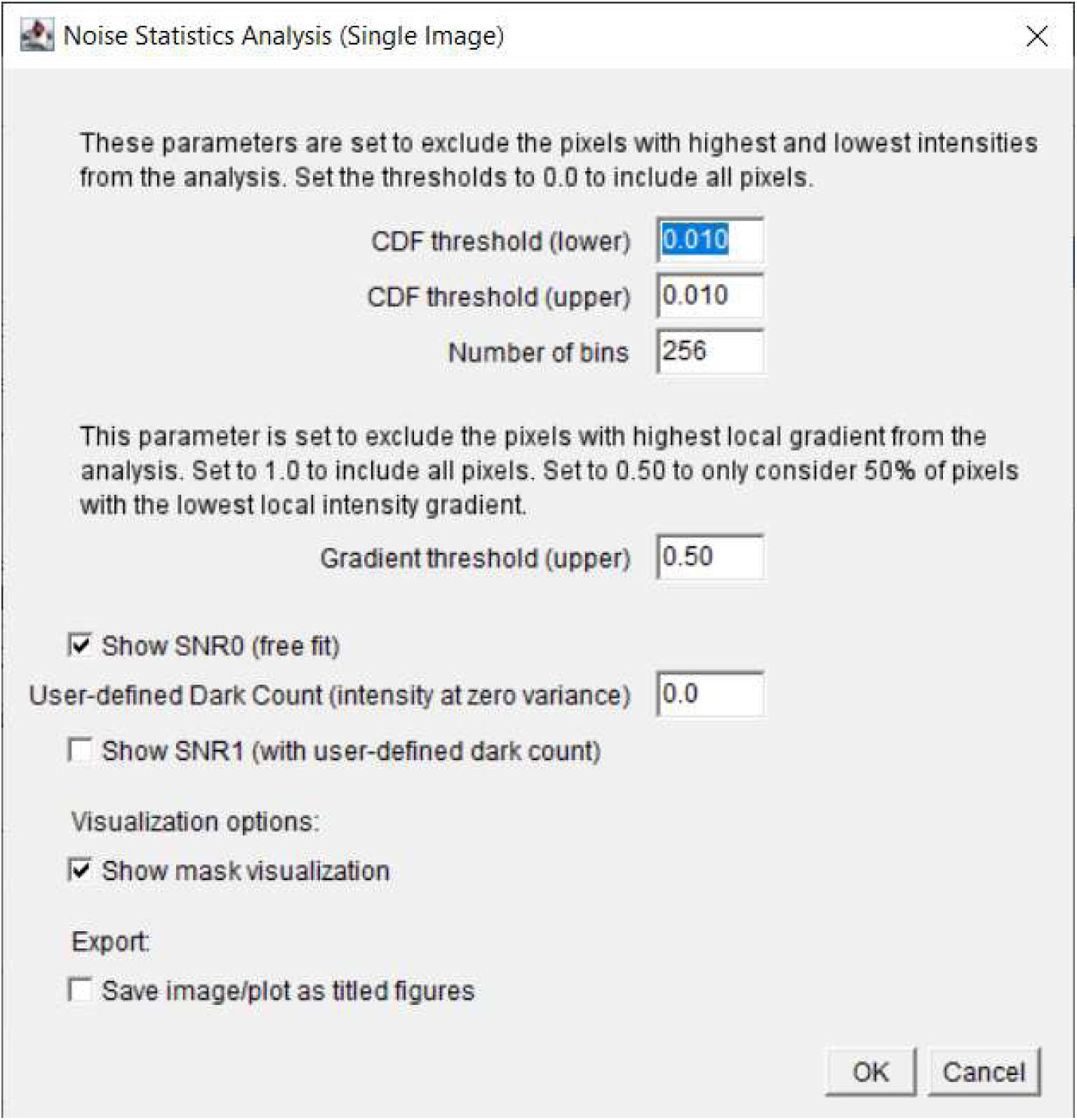
Pop-up menu with settings for Noise Analysis.

Once the plugin is executed, it will display the results as well as exclusion mask as shown in Figure 1. A separate Log window will also pop up, that will contain all settings and results in ASCII format, as shown in SI Figure 5.

**SI Figure 5.**
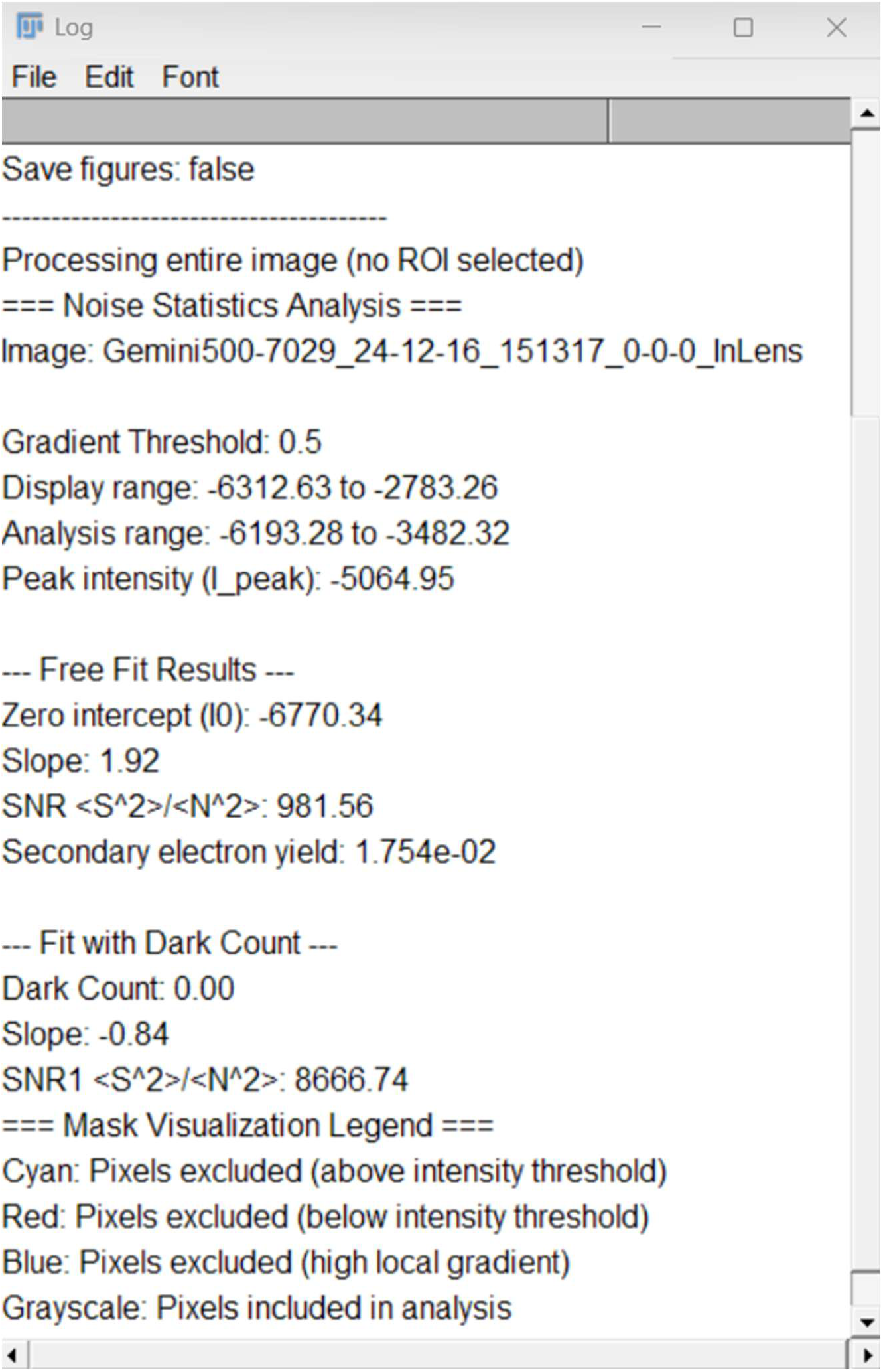
Pop-up Log window with Noise analysis results in ASCII format.

### Effect of Gradient Threshold Setting on SNR analysis results

We evaluated the effect of gradient threshold setting on SNR analysis results on images of an Sn-on-C sample imaged with 1 nm pixels (SI Figure 6, top left) and an epoxy-embedded OsO_4_-stained epithelial cell sample imaged with 4 nm pixels (SI Figure 6, top right). We varied the value of gradient threshold between 0.1 (only 10% of the pixels with lowest local intensity gradients are analyzed) and 1.0 (all pixels are analyzed) and performed SNR analysis.

The results are shown in SI Figure 6. The values of SNR (red) and Dark Count (green) as functions of the gradient threshold value are shown in SI Figure 6 (second row). The examples of the SNR analysis for the gradient threshold values of 1.0 and 0.5 are shown in SI Figure 6, third row and bottom row, respectively.

**SI Figure 6.**
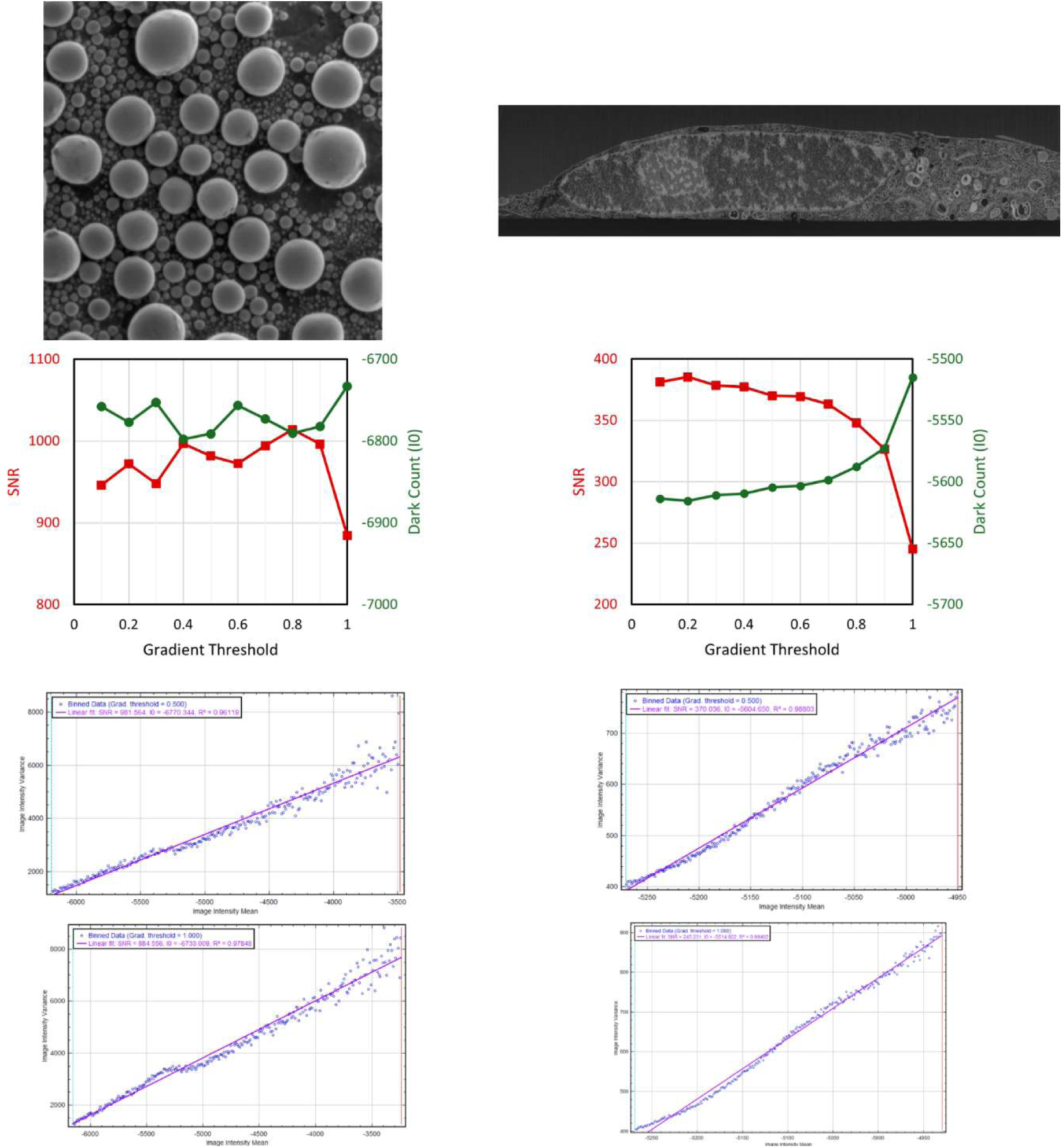
Effect of Gradient Threshold Settings on SNR analysis results.

In both cases when all pixels are included in the analysis, the Variance vs Intensity curve is not straight because the intensity variance at points with high local gradients is higher than it would be if it were determined by only shot noise. As the value of the gradient threshold decreases, the Variance vs Intensity curve becomes more linear, and the values of SNR and Dark Count stop varying. At very low values of gradient threshold, the majority of the pixels are excluded, and the analysis becomes less accurate. The setting of the gradient threshold between 0.25 and 0.50 provides the most accurate results.

### Performing the Contrast Analysis

In order to determine the contrast using Eq. (7), we need to know the value of the detector dark count, 𝐼_0_. If it is not known, it will have to be entered manually, but the accuracy of the procedure may be compromised. The contrast analysis can be performed by navigating to Plugins → FIB-SEM → Calculate Contrast. The pop-up menu with settings for Contrast analysis will appear as illustrated in SI Figure 7. In addition, the menu appears with a “live preview window,” in which you can see tints that correspond to exclusion gradient threshold and bands about the I*_low_* and I*_high_* values.

The parameters are:

**I0 handling (run SNR beforehand/manually input I0)** – if SNR wasn’t run beforehand, the user should elect whether to run the “Noise Statistics Analysis” plugin first and load the computed I0 value or to manually enter an I0 value.

**I0 (not pre-computed/pre-computed)** – detector dark count. The value will be auto-filled if the SNR analysis has been executed.

**Gradient threshold** – upper boundary for the gradient threshold. This parameter is set to exclude the pixels with high local gradients, which may help in excluding transitional areas, thus helping to isolate the high- and low-intensity peaks in PDF. The default is 0.25, meaning that only 25% of the pixels with the lowest local intensity gradients are considered in contrast analysis. There is an option to display the gradient threshold exclusion tinting if desired as well.

**I low** – adjust this parameter to identify the low-intensity peak in the PDF.

**I high** – adjust this parameter to identify the high-intensity peak in the PDF.

**I*_low_* and I*_high_* determination (auto/manual)** – check/uncheck to use a double Gaussian fitting to determine the I*_low_* and I*_high_* values or to enter those values manually.

**Plot Gaussian components (for auto only)** – if using a double-Gaussian fit, check/uncheck to display the two individual Gaussian curves and a double-Gaussian curve with an R^2^ value (goodness-of-fit measure w.r.t. the PDF).

**Number of bins (analysis)** – number of bins used for analysis of the data. Default is 256.

**SI Figure 7.**
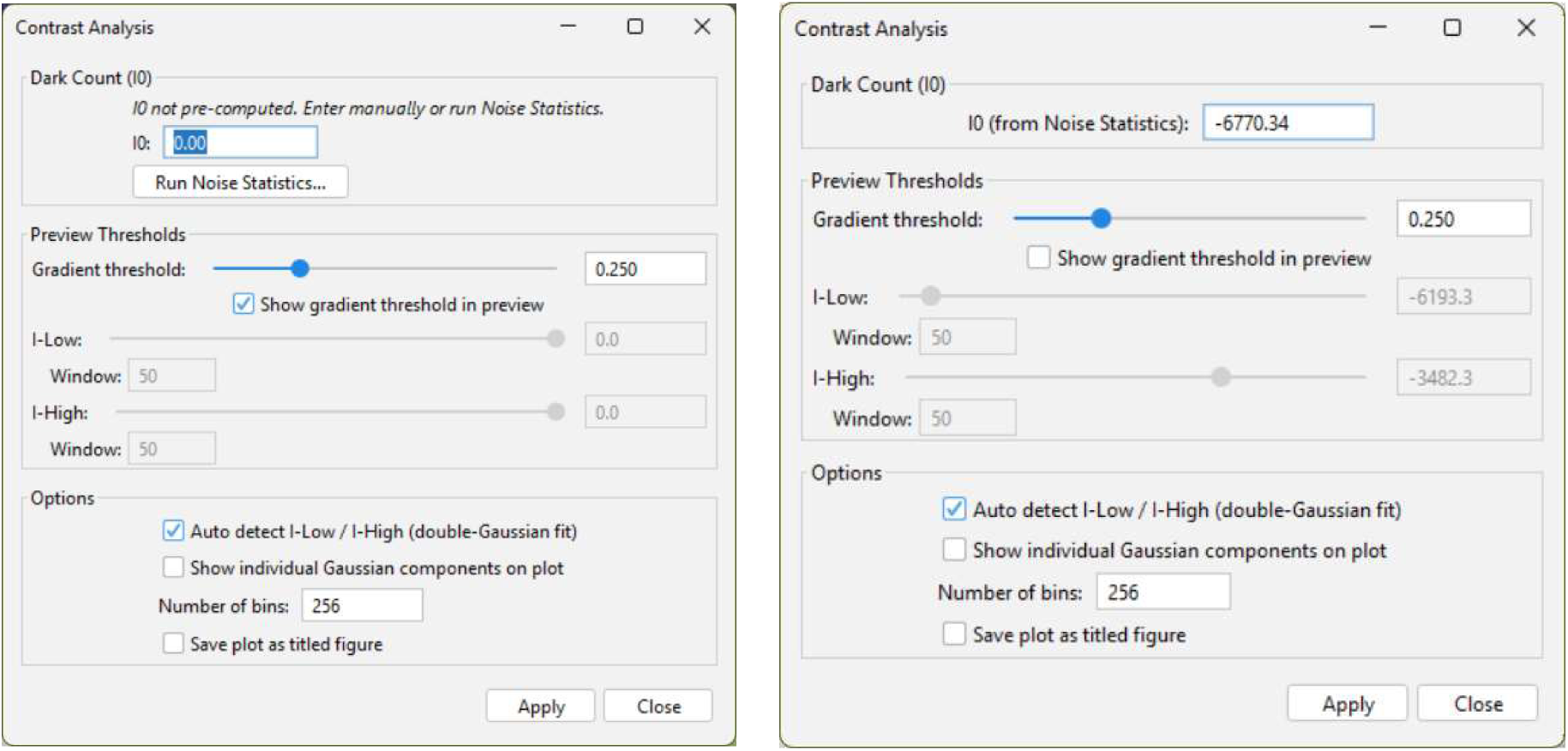
Pop-up menu with settings for Contrast Analysis. If the plugin is called before running an SNR analysis, the left dialog will appear; otherwise, the right dialog will appear.

Once the plugin is executed, it will display the results. A separate Log window will also pop up containing all settings and results in ASCII format, as shown in SI Figure 8.

**SI Figure 8.**
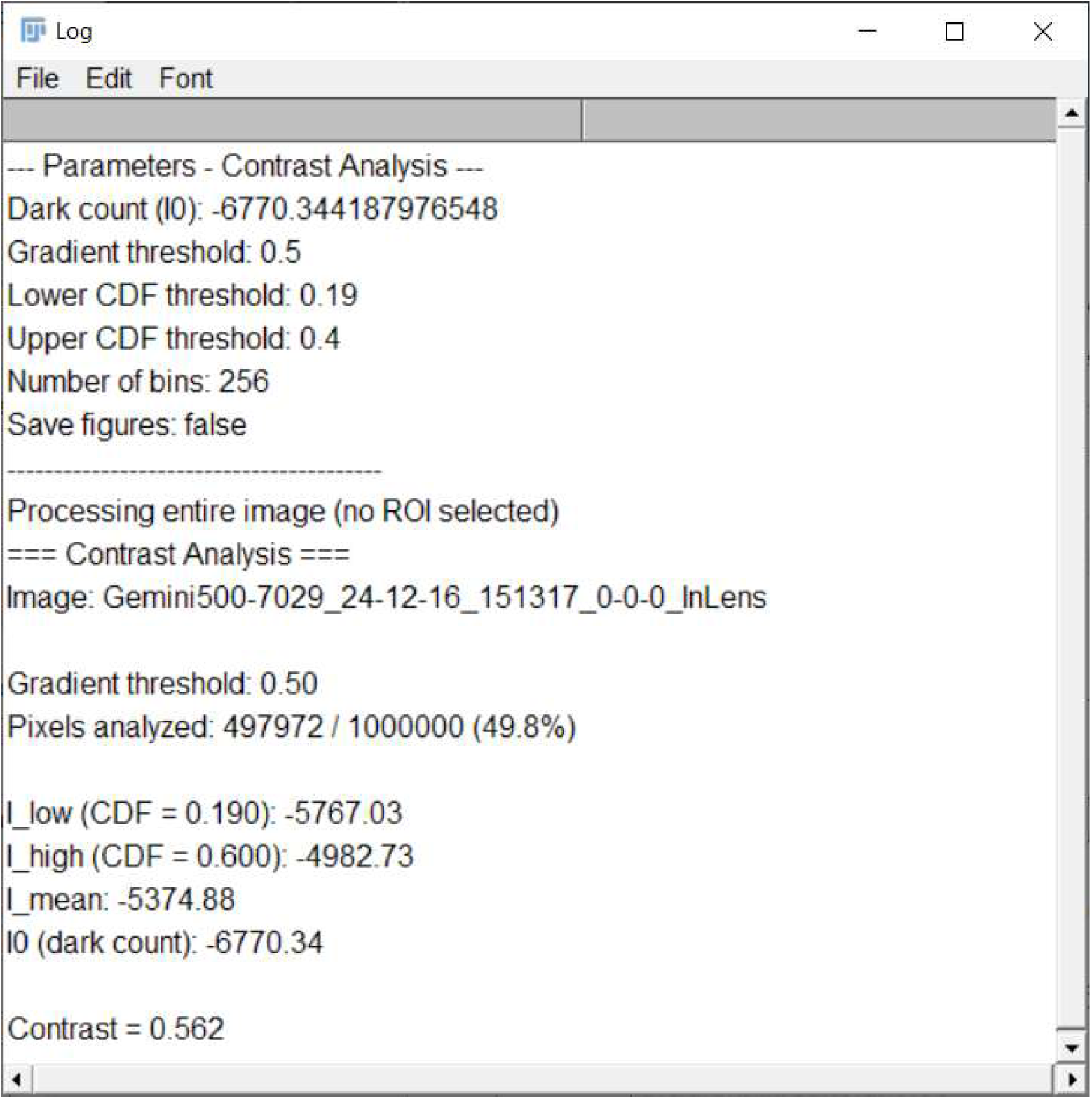
Pop-up Log window with Contrast analysis results in ASCII format.

### Performing Edge Transition Analysis

To perform the Edge Transition (resolution) analysis, navigate to Plugins → FIB-SEM → Resolution: Edge Transitions. The pop-up menu with settings for edge Transition Analysis will appear as illustrated in SI Figure 9.

The parameters are:

**Lower bound** – the lower value (start point of transition) for intensity change analysis across a sharp edge. Default is 0.37.

**Upper bound** – the higher value (end point of transition) for intensity change analysis across a sharp edge. Default is 0.63.

**Pixel Size** – pixel size in nm. If the image is imported from a .dat file, this value will be auto-populated from the header.

**Subset size** – linear size (in pixels) of the square subset selected for each point with high local gradient identified for analysis. Default is 25.

**Section length** – length (in pixels) of the intensity profile extracted for each point with a high local gradient and used for transition analysis. It cannot be larger than the subset size. Default is 25.

**Min/max aperture** – the lengths of the subsets of the intensity profile (determined above) used to calculate the minimum and maximum values of the trace.

**Gradient Threshold** – the cumulative threshold used to select the points with the highest absolute value of the local intensity gradients. At the first step, the gradient map is calculated, and a list of potential transition points is created. Only a fraction of the points, defined by the Gradient Threshold value, are retained for the subsequent analysis.

**Minimum threshold criterion** – only the points with minimum value of trace within this fraction from the overall image minimum will be analyzed.

**Maximum threshold criterion** – only the points with the maximum value of trace within this fraction from the overall image maximum will be analyzed.

**Neighbor exclusion radius** (pixels) – as transition points are selected in order of local intensity gradient, the exclusion circles are drawn for each point to exclude close neighbors.

**Center exclusion radius** (pixels) – useful if the image has a center mark (present in some EM images)

**Transition low limit** – ignore results with transition distances below this.

**Transition high limit** – ignore results with transition distances above this.

**SI Figure 9.**
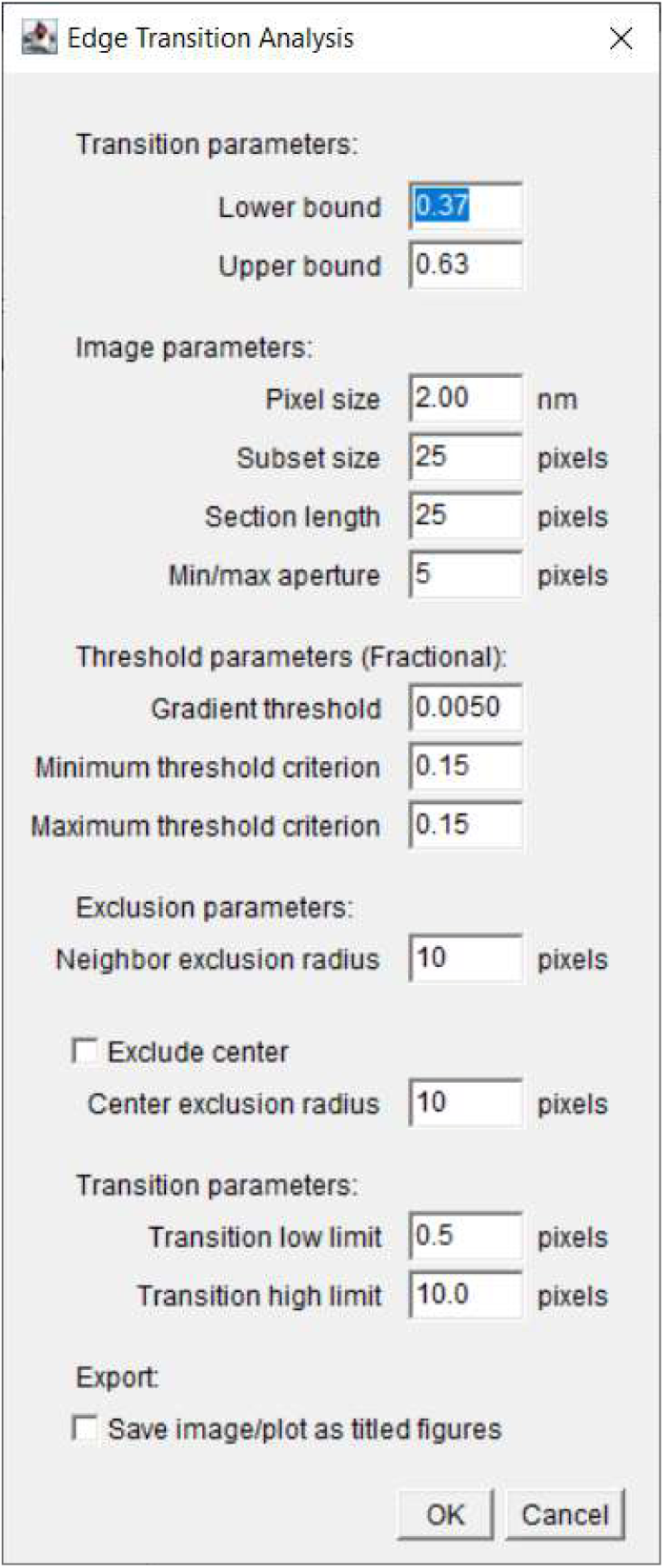
Pop-up menu with settings for Edge Transition Analysis.

Once the plugin is executed, it will display the results as shown in Figure 3. A separate Log window will also pop up containing all settings and results in ASCII format, as shown in SI Figure 10.

**SI Figure 10.**
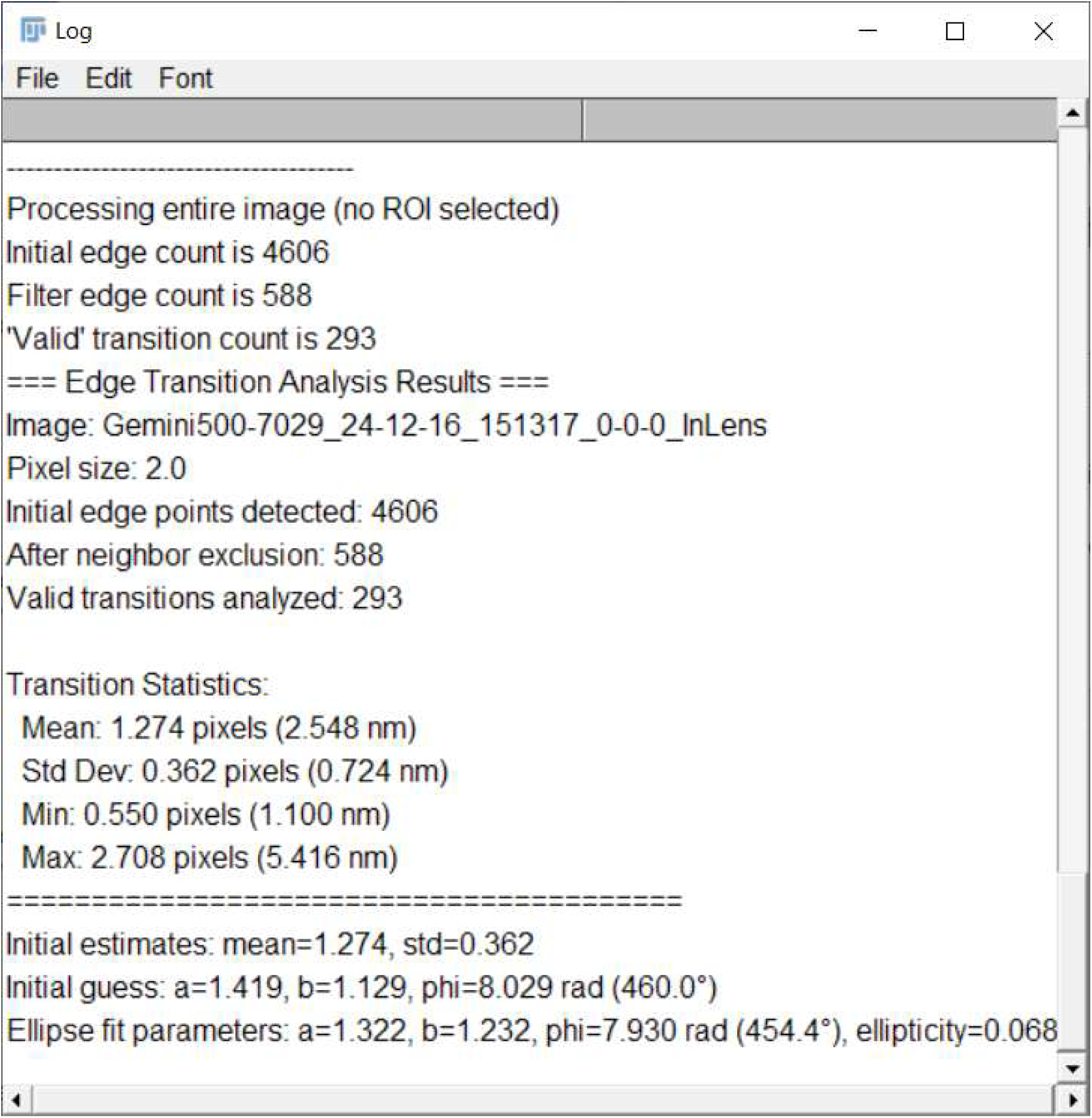
Pop-up Log window with Edge Transition analysis results in ASCII format.

## Results

### Mouse dorsal striatum

**SI Figure 11.**
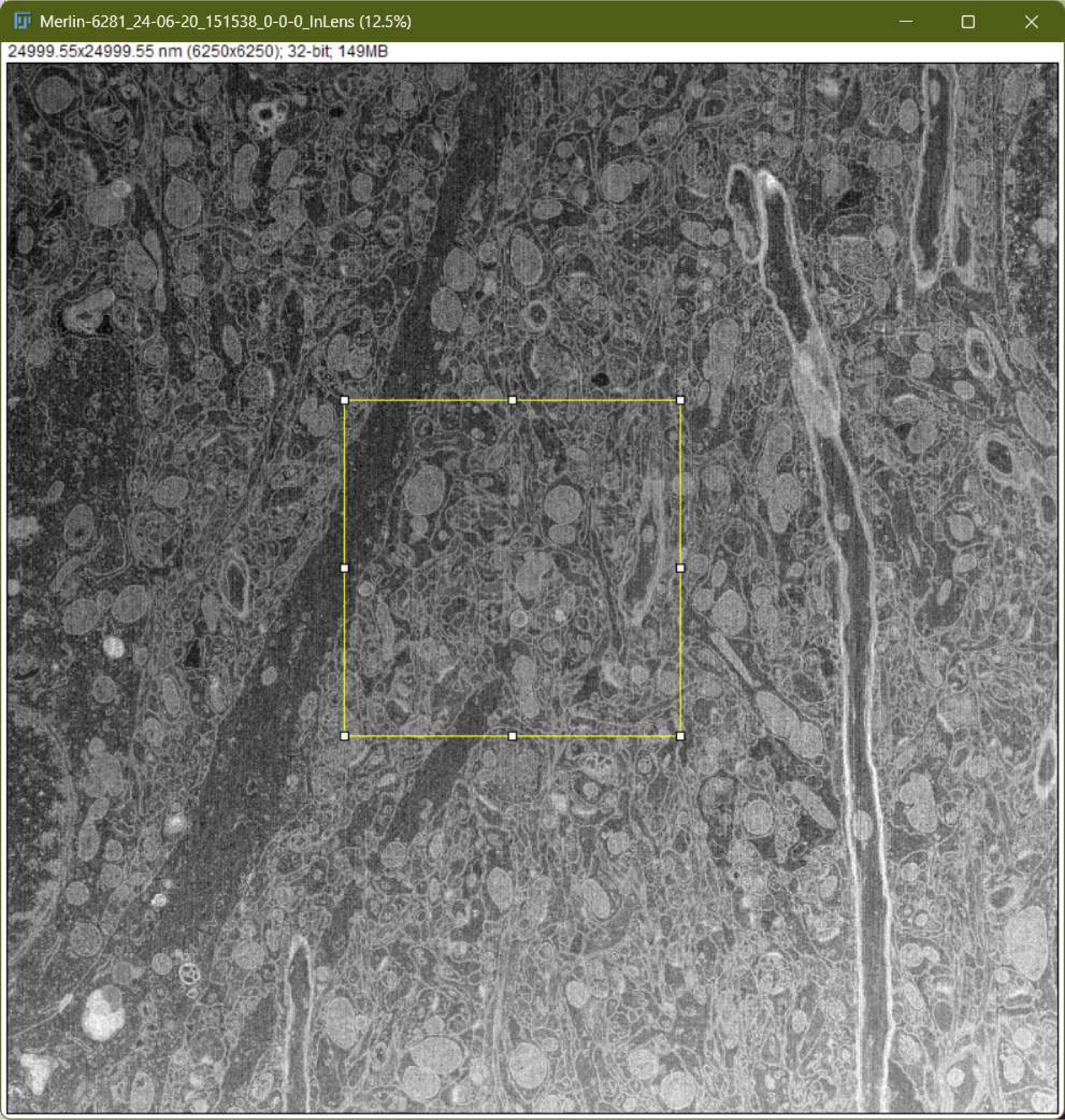
Secondary electron detector image (single frame) of the Mouse dorsal striatum sample (doi.org/10.25378/janelia.28754705).

**SI Figure 12.**
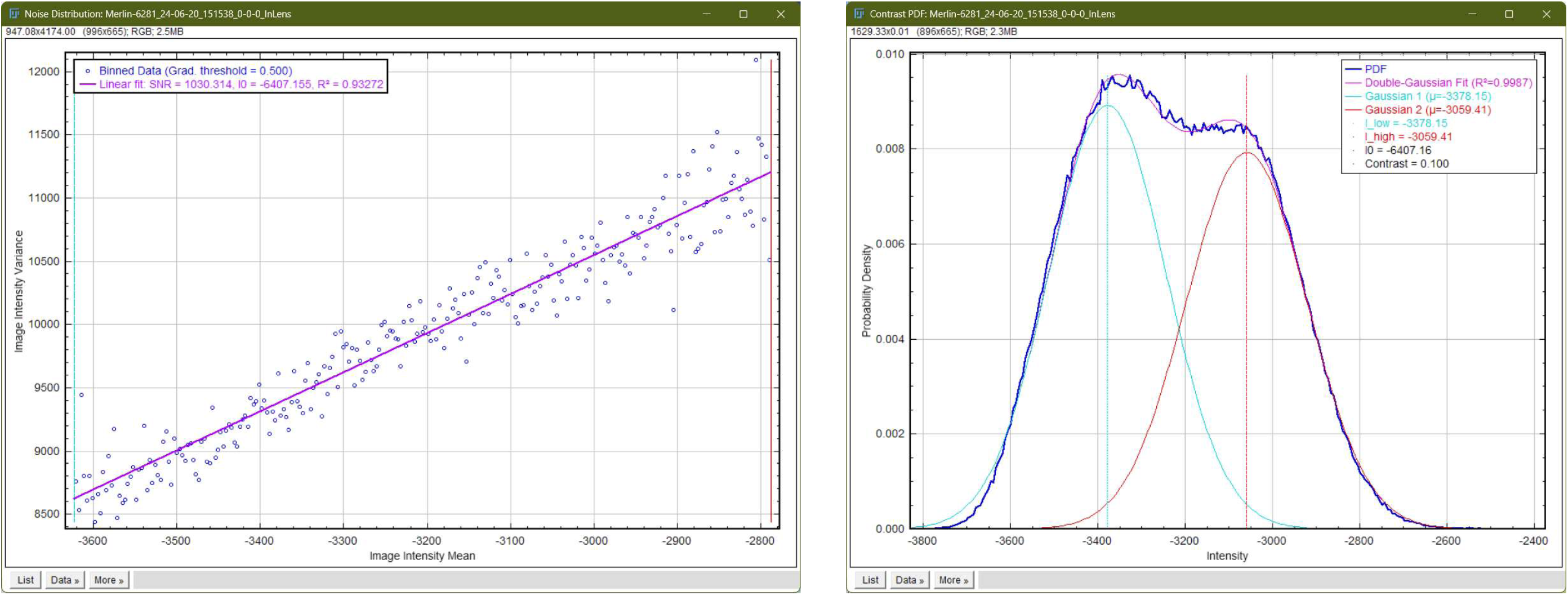
SNR and Image contrast analysis performed on a subset of a single frame of the Mouse dorsal striatum sample in SI Figure 11 (doi.org/10.25378/janelia.28754705).

### Killer T-Cell attacking cancer cell

**SI Figure 13.**
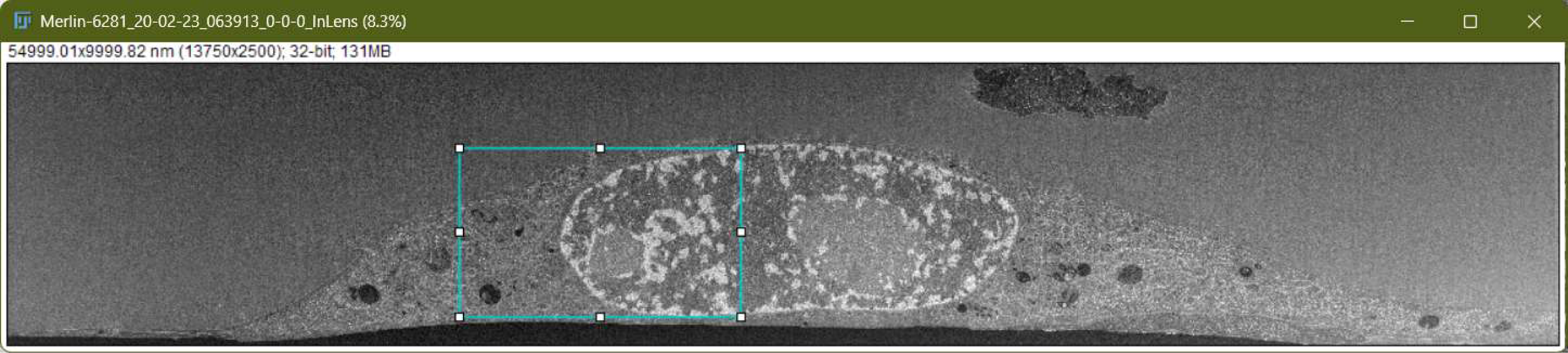
Secondary electron detector image (single frame) of the Killer T-Cell attacking cancer cell sample (doi.org/10.25378/janelia.13114454).

**SI Figure 14.**
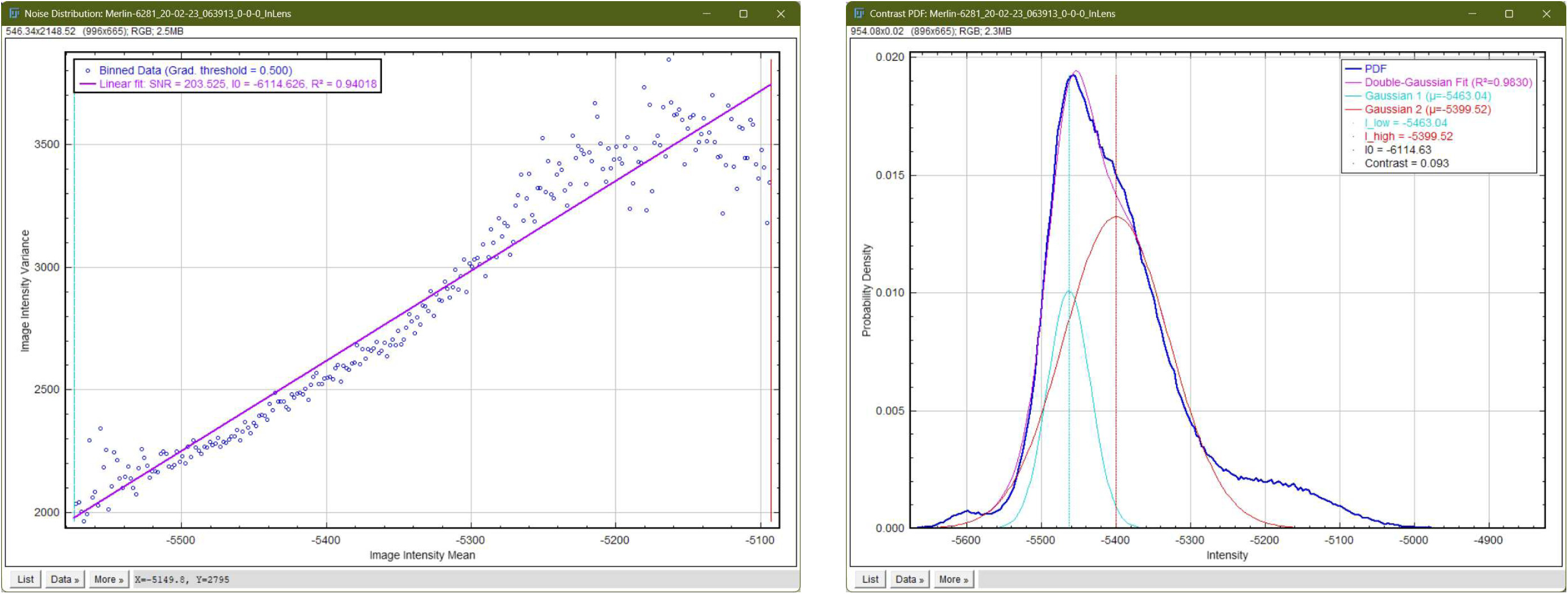
SNR and Image contrast analysis performed on a subset1 (indicated by a cyan ROI in SI Figure 13) of a single frame of the Killer T-Cell attacking cancer cell sample (doi.org/10.25378/janelia.13114454).

**SI Figure 15.**
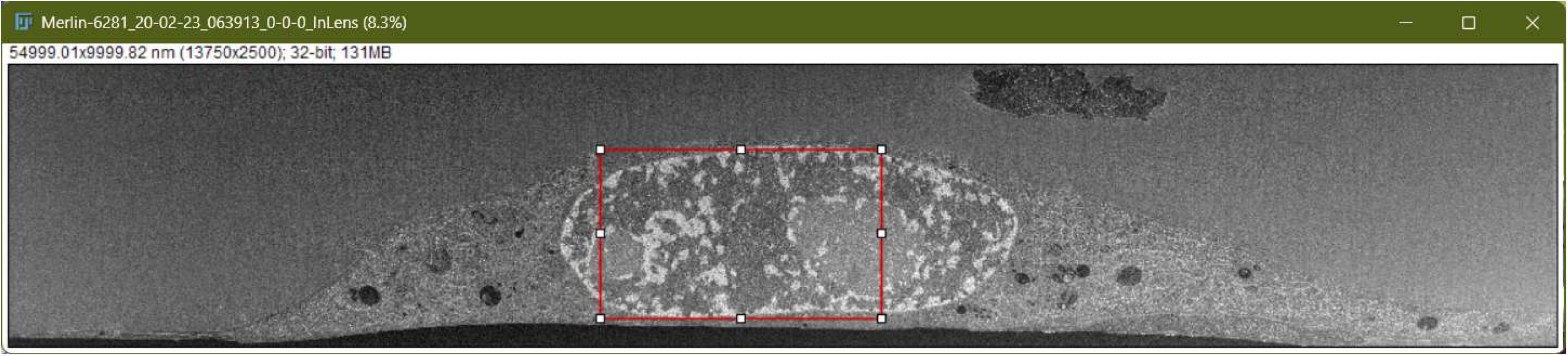
Secondary electron detector image (single frame) of the Killer T-Cell attacking cancer cell sample (doi.org/10.25378/janelia.13114454).

**SI Figure 16.**
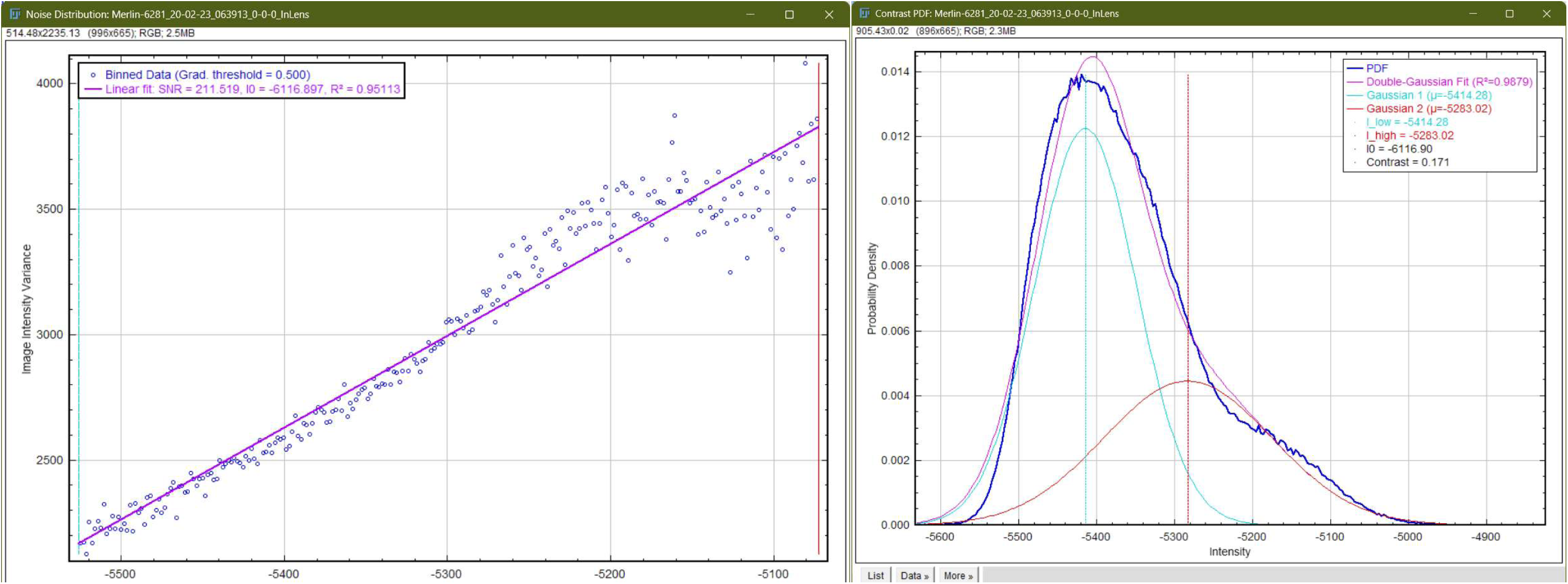
SNR and Image contrast analysis performed on a subset2 (indicated by a red ROI in SI Figure 15)SI Figure 13 of a single frame of the Killer T-Cell attacking cancer cell sample (doi.org/10.25378/janelia.13114454).

**SI Figure 17.**
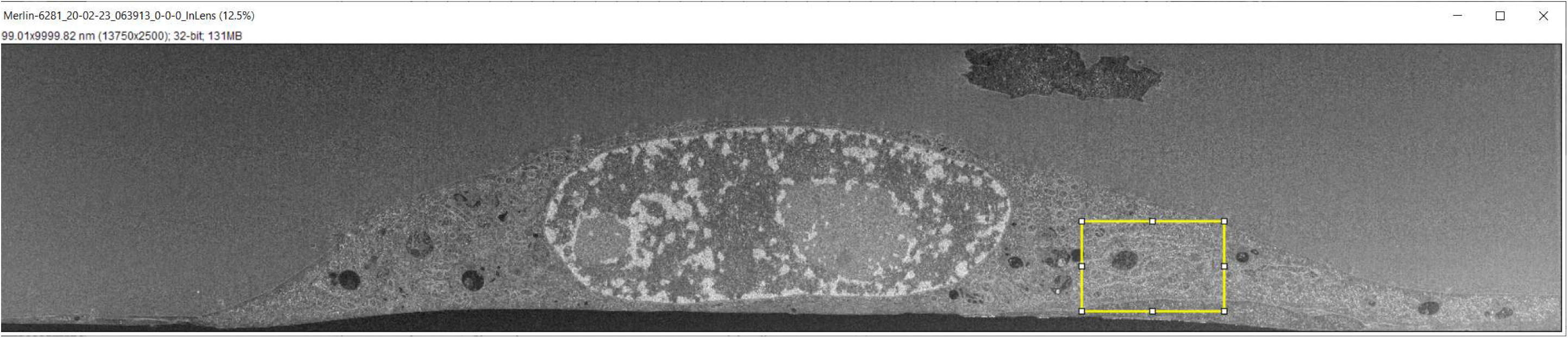
Secondary electron detector image (single frame) of the Killer T-Cell attacking cancer cell sample (doi.org/10.25378/janelia.13114454).

**SI Figure 18.**
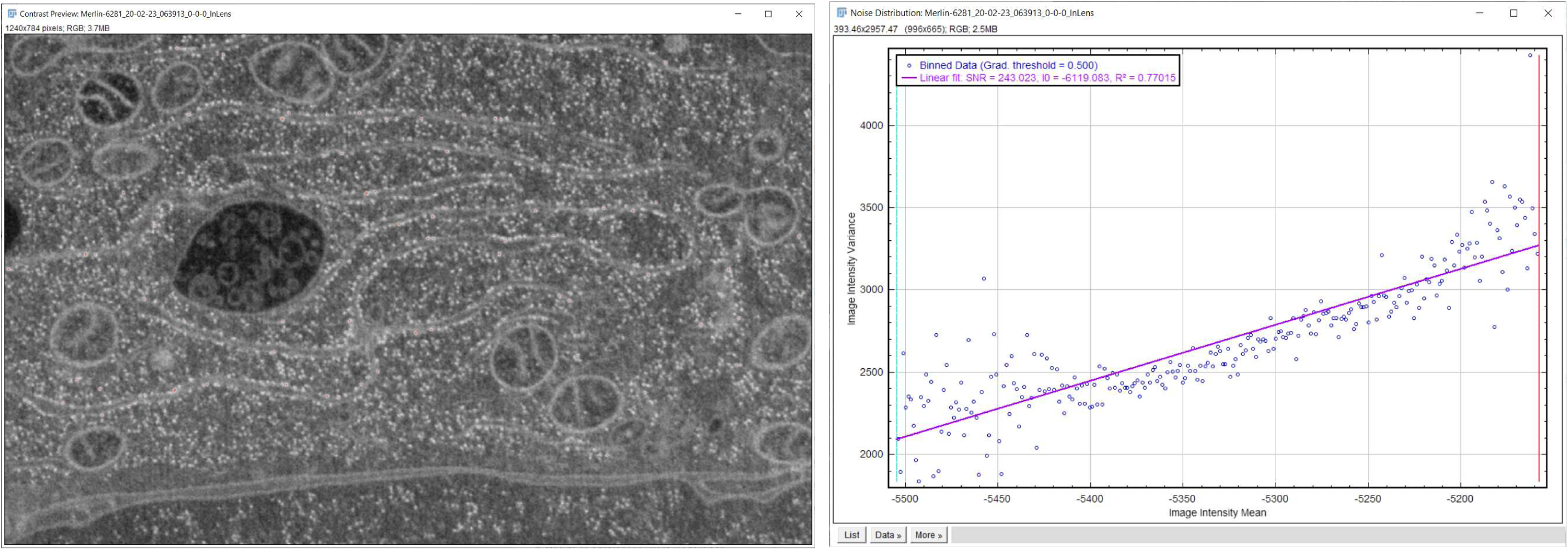
Zoomed-in section of the image in SI Figure 17, indicated by yellow rectangle (left), and results of SNR analysis performed on that section (right).

**SI Table 1.**
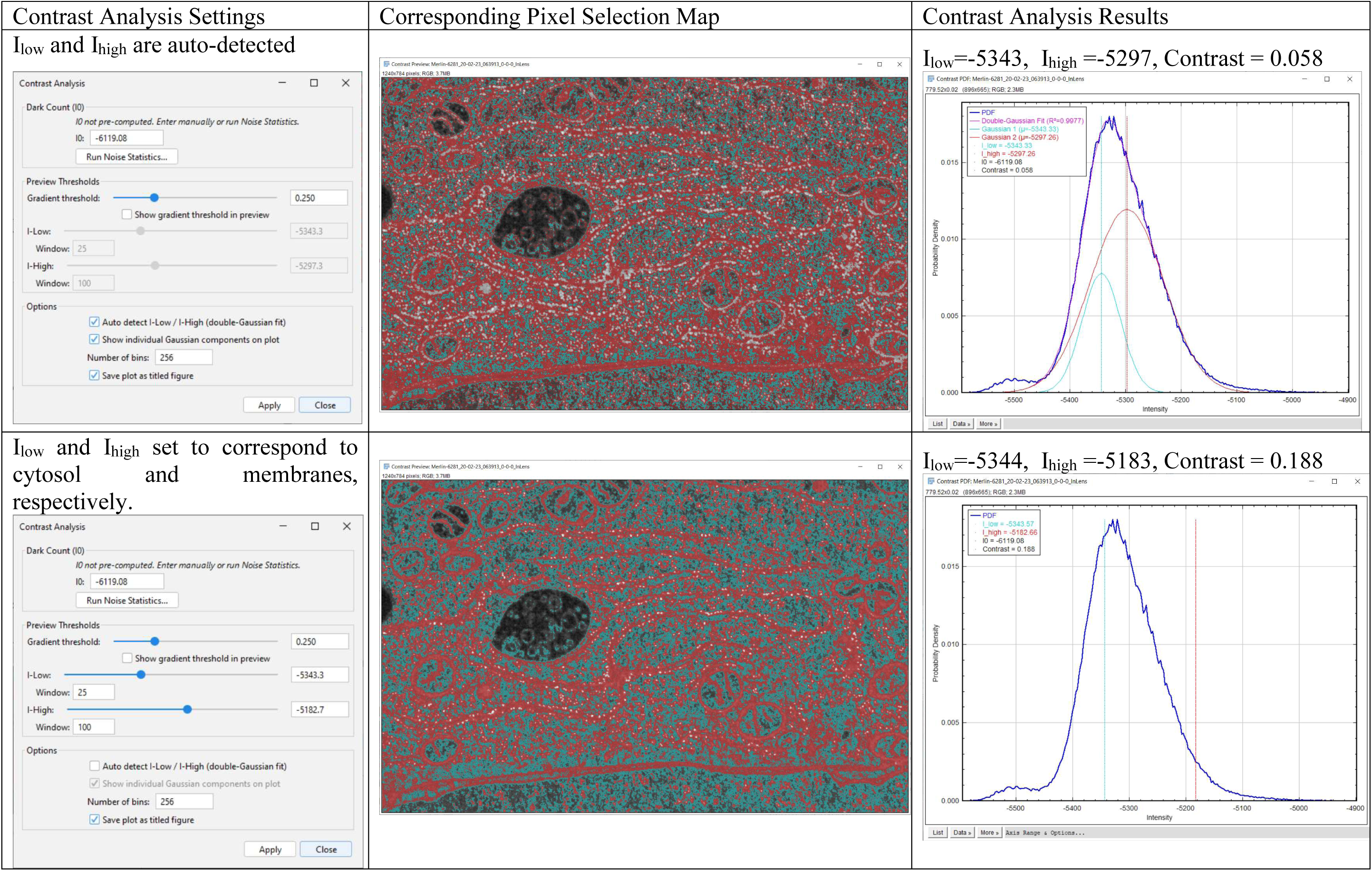
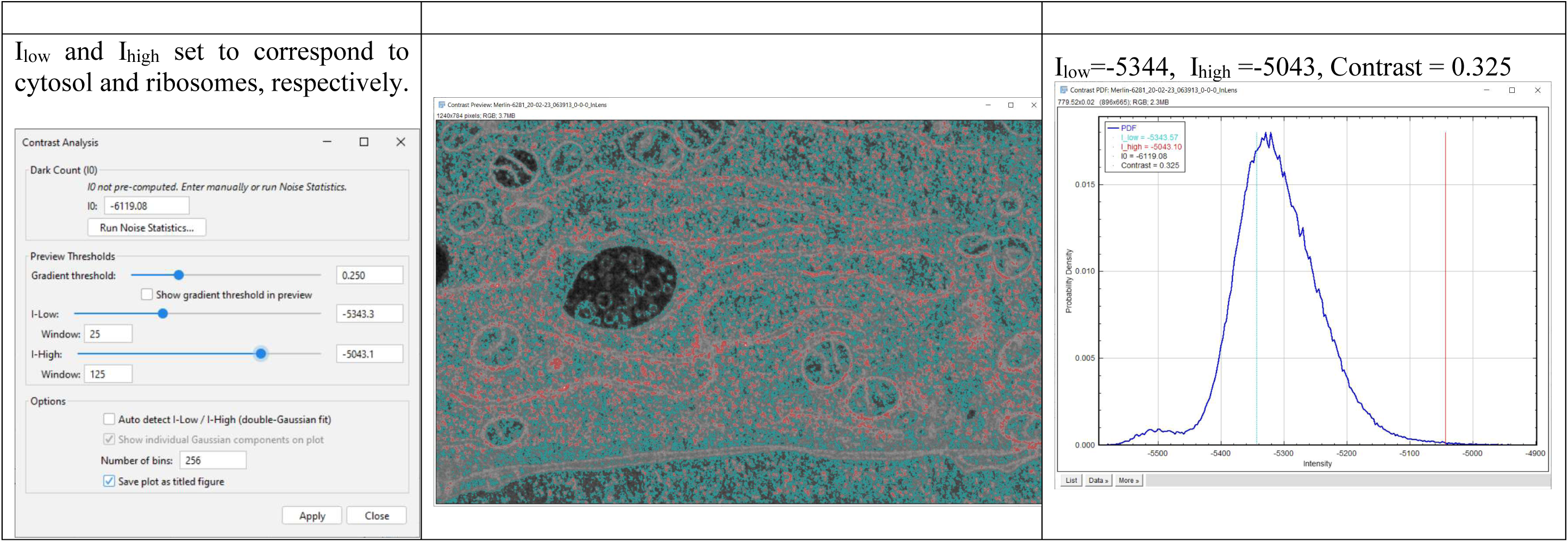
Contrast determined with auto-detected of Ilow and Ihigh levels (top row), and with Ilow and Ihigh levels manually set to correspond to cytosol and membranes, respectively (middle row), and with Ilow and Ihigh levels manually set to correspond to cytosol and ribosomes, respectively (bottom row).

### Isolated murine pancreatic islets

**SI Figure 19.**
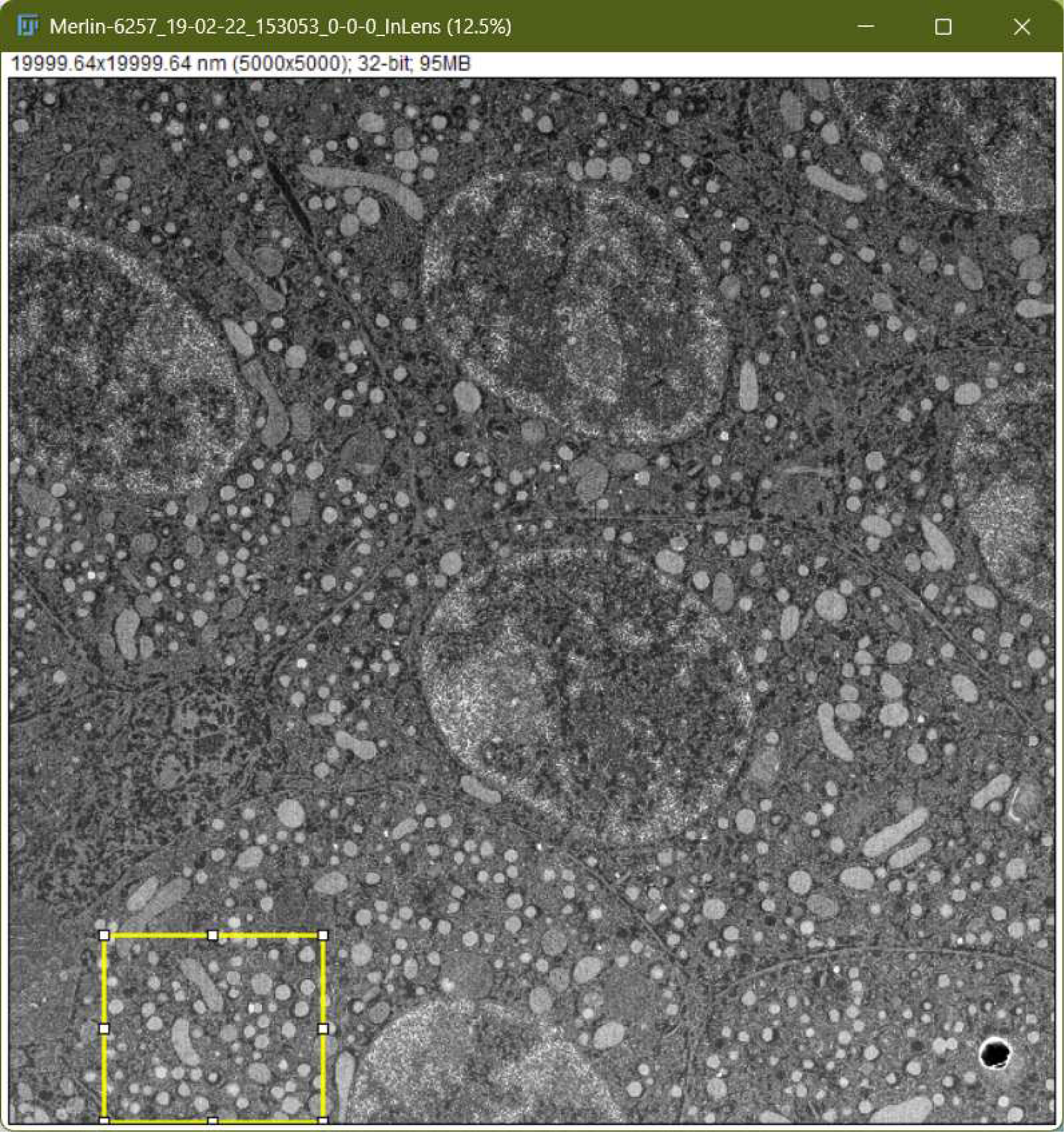
Secondary electron detector image (single frame) of the Isolated murine pancreatic islets sample (doi.org/10.25378/janelia.19196540).

**SI Figure 20.**
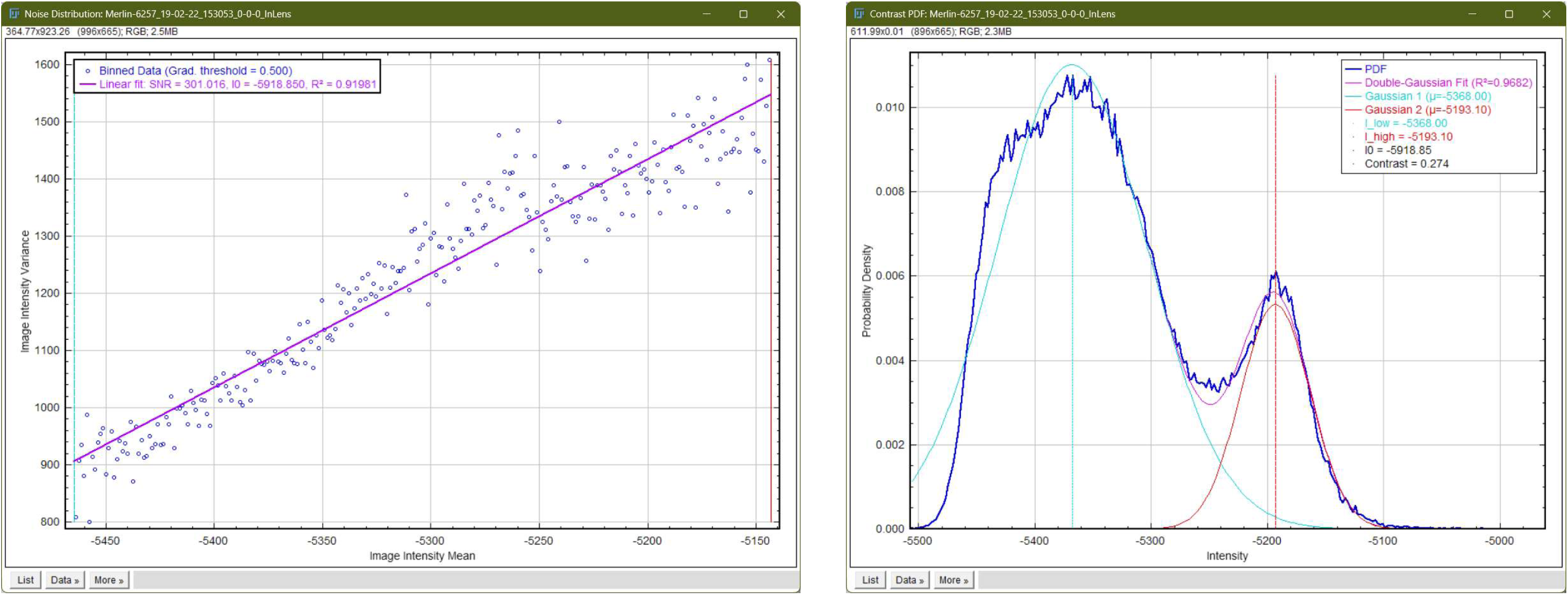
SNR and Image contrast analysis performed on a subset of Isolated murine pancreatic islets sample (doi.org/10.25378/janelia.19196540).

### Caenorhabditis elegans exposed to 60 nm Au nanoparticles

**SI Figure 21.**
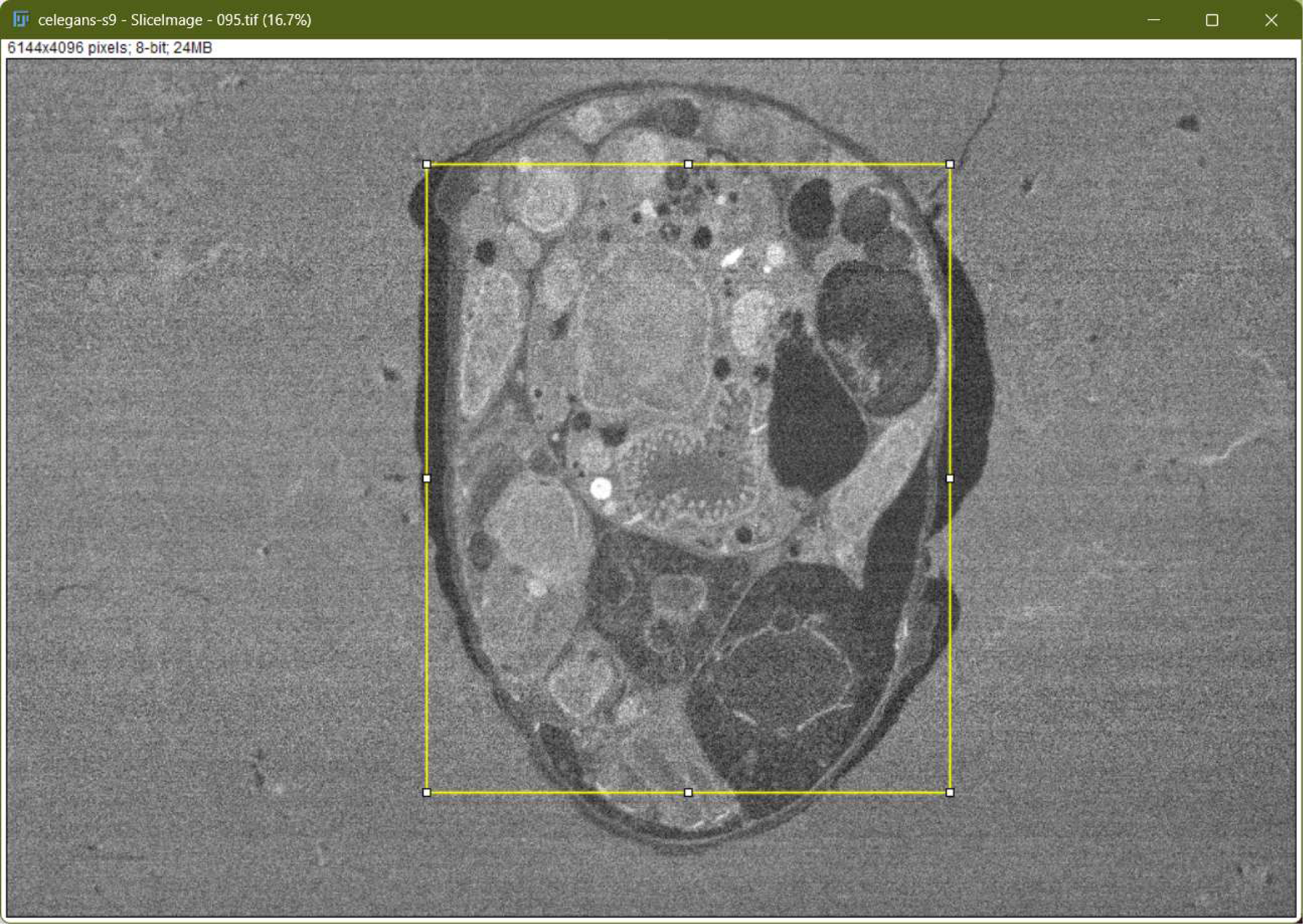
Back-scattered electron detector image (single frame) of the C. elegans sample (doi.org/10.18434/M3C09F).

**SI Figure 22.**
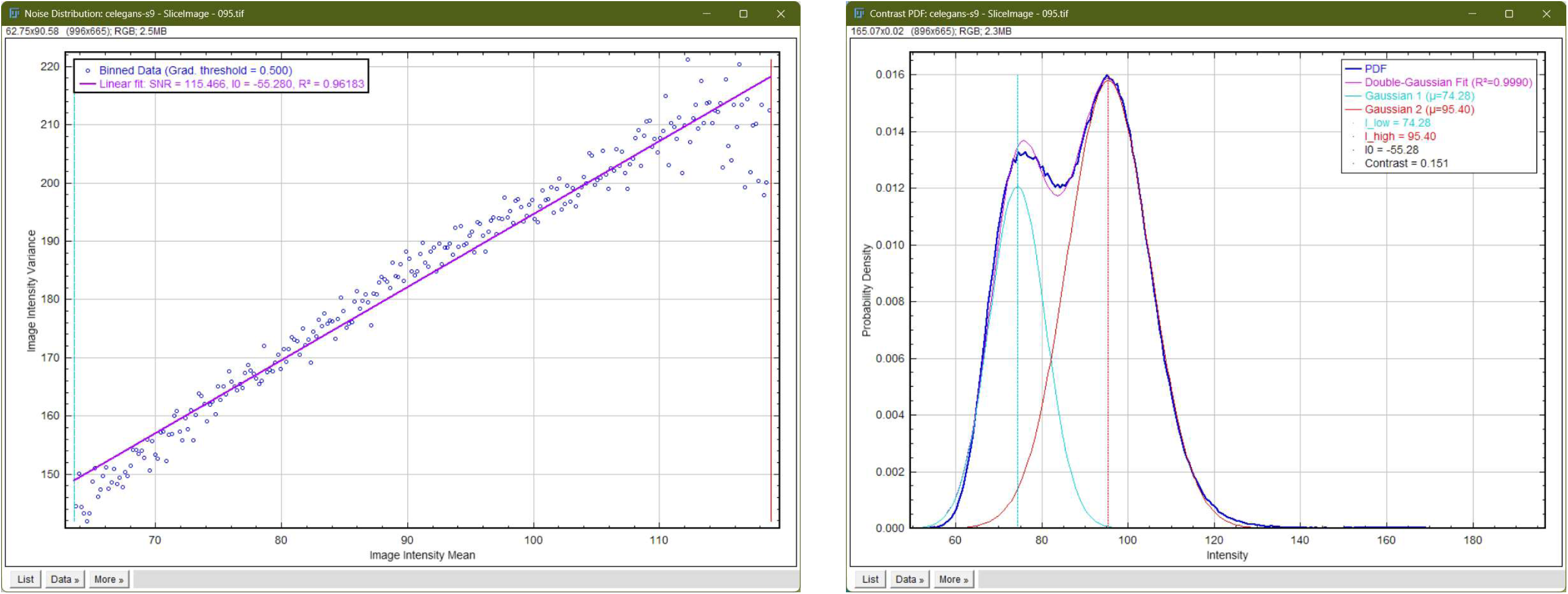
SNR and Image contrast analysis performed on a subset of *C. elegans* sample (doi.org/10.18434/M3C09F).

### Source Code

Fiji plugin is written in Java. The source code is available here: https://github.com/davidshtengel/fibsem-fiji

The Python source code is available here: https://github.com/gleb-shtengel/FIBSEM_gs_py

